# Endothelial protein C receptor is an entry receptor for Langya and related parahenipaviruses

**DOI:** 10.64898/2026.11.01.763137

**Authors:** Zhe Zhao, Paul Tran, Ning Li, Li Ou, Jaewon Song, Michael Sibilo, Mike Castro, Ananda Chowdhury, Jennifer A. Bohl, Jason M. Wu, Thythy T. Do, Baoshan Zhang, Jenny Zhao, Anasuya C. Pal, Clancy W. Mullan, David R. Martinez, Richard W. Pierce, Alicen Spaulding, Sanghyun Lee, Yaroslav Tsybovsky, Leonid Serebryannyy, Daniel C. Douek, Tongqing Zhou, Craig B. Wilen

## Abstract

Langya virus is a zoonotic parahenipavirus with pandemic potential that causes acute febrile illness in people. We identified endothelial protein C receptor (EPCR) as an entry receptor for Langya, Wenzhou, and Mojiang parahenipaviruses. A 3.1 Å cryo-electron microscopy structure reveals that Langya attachment glycoprotein engages EPCR through a site distinct from that of Nipah’s ephrin binding site. This binding mode resembles those of protein C and a malarial adhesin, demonstrating convergent evolution. Interface substitutions abolish entry, whereas natural variation explains differential human EPCR use and predicts receptor compatibility across 26 animal species. These findings establish EPCR as a receptor for the Langya clade and provide a structural framework for understanding cross-species transmission and human infection potential.

## Main text

Parahenipaviruses are an emerging group of viruses in the Paramyxoviridae family that pose a significant yet poorly understood threat to biosecurity and public health. Parahenipaviruses are related to henipaviruses, including highly pathogenic Nipah and Hendra viruses, which can cause acute respiratory distress syndrome (ARDS) and encephalitis with a 40-75% mortality rate(*1*, *2*). While fruit bats in Asia and Australia are the primary reservoirs of Nipah and Hendra, parahenipaviruses have been detected in multiple shrew, rodent, and bat species across multiple continents, including North America(*3–8*). The clinical relevance of parahenipaviruses was demonstrated by the initial discovery of Langya virus in 35 patients with febrile illness, leukopenia, and organ dysfunction(*3*). Langya virus was subsequently identified in two shrew species (*Crocidura lasiura* and *Crocidura shantungensis*). Inadequate surveillance and diagnostic testing limit the accurate assessment of human disease burden(*6*). The related Mojiang virus was identified in rats (*Rattus tanezumi*) during an investigation of fatal pneumonia in three miners who cleared bat guano from a mine shaft in Mojiang, China(*5*). However, a causal link for Mojiang has not been established. Wenzhou virus, which is in the same parahenipavirus clade as Langya and Mojiang viruses, has been detected in both the Asian house shrew (*Suncus murinus*) and striped field mouse (*Apodemus agrarius*), but its ability to infect people and cause disease remains unknown^4^.

Henipa- and parahenipavirus entry requires the coordinated action of the viral attachment glycoprotein (G) and fusion glycoprotein (F). G forms a tetramer, with each head domain adopting a six-bladed β-propeller fold that engages the host receptor(*9*, *10*). The fusion glycoprotein (F) is a class I trimer that is activated by endosomal cathepsin cleavage(*11*). G and F associate on the virion surface, and receptor engagement by G releases F to insert its fusion peptide into the target membrane which drives membrane fusion(*11*),(*12*). While Nipah and Hendra viruses use ephrin-B2 and ephrin-B3 as receptors, parahenipavirus receptors are unknown(*13–16*). This gap has limited our understanding of parahenipavirus cell and tissue tropism, host range, and zoonotic potential, while hindering efforts to culture infectious viruses, establish animal models, and develop countermeasures, including neutralizing antibodies and vaccines.

### Human EPCR is sufficient for Langya binding and entry

To define the evolutionary relationships among henipa- and parahenipaviruses, we inferred a maximum-likelihood phylogeny of the viral G protein and annotated each virus by reservoir host and country of detection (**Fig. 1A**). Parahenipaviruses formed a clade distinct from henipaviruses. Within this clade, Langya, Wenzhou, and Mojiang viruses clustered together. The Langya G ectodomain sequenced from patients is identical to that recovered from *Crocidura lasiura* shrews in the same survey in China and 97% identical to more recently sampled sequences from South Korean shrews (e.g. Langya Cl17-6)(*3*, *17*). Wenzhou G ectodomain sequences from the Asian house shrew and striped field mice are 98% identical. For clarity, here we use the terms Langya and Wenzhou to refer to viral sequences from human and shrew (*Suncus murinus*), respectively.

**Figure 1.**
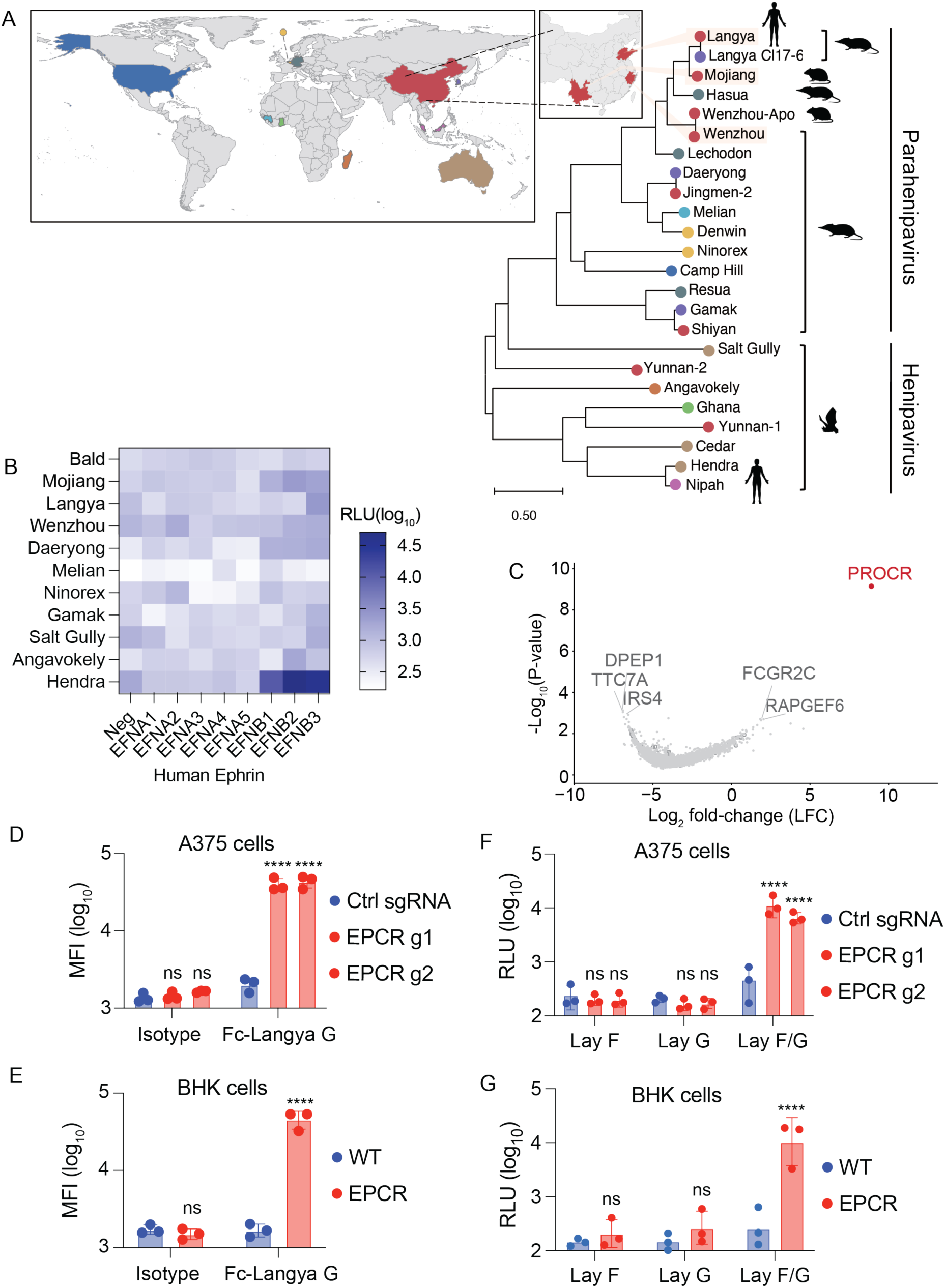
Human EPCR is sufficient for Langya binding and entry. (**A**) Maximum-likelihood phylogeny of henipa-and parahenipavirus attachment glycoprotein, with tip color denoting country of detection (world map and China inset, left). Silhouettes denote reservoir animal host group and documented human infection. Scale bar, 0.5 substitutions per site. (**B**) HeLa-USU cells were transfected with indicated human ephrins (EFN) or a no plasmid control (neg) and challenged with VSVΔG-luc coated with G and F proteins from the indicated virus. VSVΔG-luc without G/F (bald) and with Hendra G/F are included as negative and positive controls, respectively. (**C**) CRISPRa screen in human A375 cells stained and sorted for Fc-Langya G binding reveals PROCR which encodes EPCR as top hit. (**D-G**) EPCR was induced in A375 cells by CRISPRa using two sgRNAs (g1, g2) (**D, F**), and expressed in hamster BHK cells by lentivirus (**E, G**). (**D, E**) Cell-surface binding of an Fc-Langya G or an isotype control, detected with anti-Fc-FITC secondary antibody and reported as mean fluorescence intensity (MFI) by flow cytometry. (**F, G**) The same cells were challenged with VSVΔG-luc coated with Langya (Lay) F alone, Langya G alone, or both G and F. Viral entry was quantified as luciferase activity (RLU, relative light units). Data in (**B**) represents the mean of each of three independent experiments each done in triplicate. Each dot in (**D-G**) represents the mean of one independent experiment each done in triplicate. Error bars, SD. Data analyzed by two-way ANOVA with Dunnett’s or Sidak’s multiple comparisons test versus control sgRNA (**D, F**) or parental BHK cells (**E, G**). ****p<0.0001 Ns, not significant.

We first tested whether diverse parahenipaviruses could use any human ephrin as a receptor. We individually expressed all eight human ephrins (ephrin-A1-5 and -B1-3) in HeLa-USU cells which are deficient in endogenous ephrins(*18*). We challenged cells with pseudoviruses containing the G and F proteins from seven parahenipaviruses (Langya, Mojiang, Wenzhou, Daeryong, Melian, Ninorex, and Gamak), two henipaviruses with unknown receptors (Salt Gully and Angavokely), Hendra as a positive control, and mock (no G/F) as a negative control(*3–5*, *19*– *23*). We expressed G/F on replication defective vesicular stomatitis virus lacking the native VSV-G and expressing Renilla luciferase (VSVΔG-luc). As expected, Hendra G/F-VSVΔG-luc pseudovirus could use ephrin-B2 and -B3 and to a lesser extent ephrin-B1 for entry. However, none of the seven parahenipaviruses or two henipaviruses with unknown receptors exhibited significant infection using any of the human ephrins suggesting these viruses use a non-ephrin receptor (**Fig. 1B**).

To identify parahenipavirus receptor candidates, we focused on Langya virus given its human infection potential(*3*). We generated an Fc-fusion protein of the Langya G ectodomain and performed a CRISPR activation (CRISPRa) screen in the human A375 cell line(*24*). The CRISPRa library contained 24,367 sgRNAs targeting 5,891 cell-surface and membrane-associated genes, plus 1,000 non-targeting controls. We stained and FACS sorted 32 million cells for binding to Fc-Langya G and compared sgRNAs abundance in the Fc-Langya G bound relative to unsorted cells. The top hit in the screen was for the gene PROCR which encodes endothelial protein C receptor (EPCR) (**Fig. 1C**).

EPCR, also known as CD201, is a type I transmembrane glycoprotein with an N-terminal MHC-like domain that contains a hydrophobic groove that binds a phospholipid(*25–27*). EPCR is highly expressed on endothelial cells where it binds and displays its physiologic ligand, protein C, to the thrombin/thrombomodulin complex(*25*). This results in cleavage and activation of protein C which functions as a potent anti-coagulant(*28*). Activated protein C also has cytoprotective effects in the endothelium(*29*)(*25*). Interestingly, the malaria protein *Plasmodium falciparum* erythrocyte membrane protein 1 (PfEMP1) also binds EPCR through a related interface from that of protein C(*30*, *31*). PfEMP1 binding to EPCR is associated with severe malaria(*30*, *31*).

We independently validated Langya G binding to EPCR by using individual CRISPR sgRNAs in human A375 cells (**Fig. 1D**) and lentiviral transduction of EPCR encoding cDNA in hamster BHK cells (**Fig. 1E**). Next, we used VSVΔG-luc coated with Langya G/F proteins to test whether EPCR is a functional receptor for Langya virus. EPCR expression in A375 cells by CRISPRa and BHK cells by lentiviral expression was sufficient to render cells susceptible to Langya G/F mediated entry (**Fig. 1F-G**). VSVΔG-luc coated with either Langya G alone or Langya F alone was unable to enter either A375 or BHK cells stably expressing human EPCR. This is consistent with the canonical and independent attachment and fusion functions of G and F.

### Human EPCR is sufficient for related parahenipaviruses Wenzhou and Mojiang

Next, we tested if human EPCR was sufficient for entry of other parahenipaviruses. We generated VSVΔG-luc coated with G/F of Langya, Wenzhou, Mojiang, Daeryong, Melian, Ninorex, Gamak, and Angavokely and challenged BHK cells stably expressing with or without human EPCR (**Fig. 2A**). Langya and Wenzhou pseudovirus exhibited EPCR dependent entry. Mojiang pseudovirus exhibited entry ∼0.5 log above background but did not reach statistical significance. No other parahenipavirus pseudovirus could use human EPCR. Next, we used a reverse pseudovirus approach in which we express viral G/F proteins on the cell surface and challenge cells with either replication defective lentivirus or VSVΔG bearing EPCR on the virion surface. Using EPCR coated lentivirus encoding GFP we confirmed that EPCR is sufficient for entry of Langya and Wenzhou and also observed significant entry of Mojiang but not Daeryong, Melian, Ninorex, Gamak, or Angavokely. Ghana henipavirus which uses ephrin-B2 as a receptor was included as a negative control(*32*, *33*) (**Fig. 2B**). Langya, Wenzhou, and Mojiang can similarly use human EPCR when expressed on VSVΔG-luc instead of a lentivirus (**Fig. 2C**). Notably,

**Figure 2.**
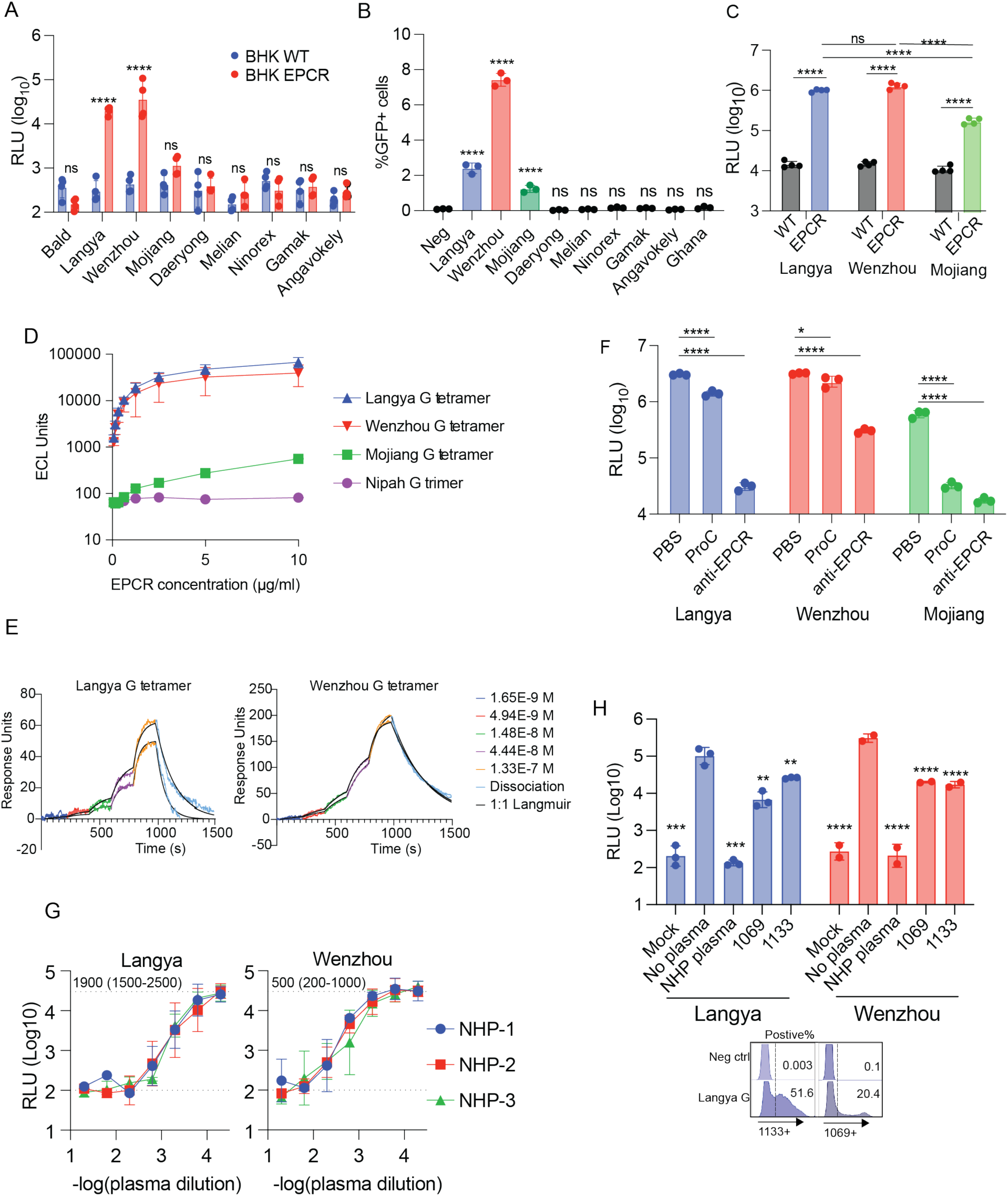
Human EPCR is sufficient for related parahenipaviruses Wenzhou and Mojiang. (**A**) Wild-type (WT) BHK cells or BHK cells stably expressing human EPCR were challenged with VSVΔG-luc coated with G/F of indicated parahenipaviruses. Entry was quantified as luciferase activity (RLU, relative light units). Bald refers to VSVΔG-luc without G/F (**B**) 293T cells were transfected with G and F of respective parahenipaviruses or a Ghana virus control. Cells were challenged with human EPCR coated lentivirus expressing GFP. Percent GFP+ cells are shown. Data is representative of one of two independent experiments performed in triplicate and is analyzed by one-way ANOVA with Dunnett’s multiple comparisons test relative to Langya F only negative control (neg). (**C**) 293T cells were transfected with G/F of Langya, Wenzhou, or Mojiang and then challenged with VSVΔG-luc coated with human EPCR. (**D**) Binding (ECL units) of varying concentrations of human ECPR to Langya G tetramer, Wenzhou G tetramer, Mojiang G tetramer, and Nipah G trimer on MSD plates. Data is representative of two experiments done in duplicate. (**E**) Langya and Wenzhou G protein binding to human EPCR as determined by single cycle kinetics SPR in two independent experiments. Apparent kinetic constants are shown as averages ± standard deviations. (**F**) VSVΔG-luc coated with human EPCR was incubated with recombinant protein C at 20 µg/ml, anti-EPCR (clone RCR-401) at 10 µg/ml or a PBS control and then added to HEK293T cells expressing parahenipavirus G/F proteins. (**G**) VSVΔG-luc coated with G/F of Langya or Wenzhou was mixed with vaccine plasma from three non-human primates (NHP-1-3) and then added to BHK cells stably expressing human EPCR cells. Geometric mean IC50 values and 95% CI are shown. (**H**) Langya G/F or Wenzhou G/F coated VSVΔG-luc was incubated with 1:10 NHP Langya vaccine plasma, or 10 µg/ml of anti-Langya clones 1069 and 1133 and added to BHK cells stably expressing human EPCR. Mock refers to VSVΔG-luc without G/F. In (**A, C, F**, **H**), entry was quantified as luciferase activity (RLU, relative light units). 1069 and 1133 binding to Langya was confirmed by flow cytometry. Data in (A, C, F-H) represents the mean+/-SD of 3-4 independent experiments each performed in triplicate except for Wenzhou in (H) which was performed in two independent experiments. Data in (**A, B, C, F, H**) is analyzed by two-way ANOVA with Dunnett’s or Sidak’s multiple comparisons test. *p<0.05, **p<0.01, *** p< 0.001, ****p<0.0001 Ns, not significant.

Langya and Wenzhou use of human EPCR is more efficient than that of Mojiang across all three assays (**Fig. 2A-C**). The reverse pseudovirus approach confirms G/F interactions with EPCR alone are sufficient for membrane fusion and suggests that no additional host surface receptor or coreceptor is required for G/F-mediated entry(*34*).

To assess binding affinity of Langya, Wenzhou, and Mojiang G to human EPCR, we titrated soluble EPCR against immobilized G tetramers by Meso Scale Discovery electrochemiluminescence (**Fig. 2D**). All three parahenipavirus G tetramers bound in a concentration-dependent manner while a negative control Nipah G trimer remained at background. The apparent Kd values were 59 +/-19 nM for Langya, 31 +/- 20 nM for Wenzhou, and 260 +/- 160 nM for Mojiang. The Mojiang apparent Kd exceeds the highest concentration tested and therefore represents a lower bound. Surface plasmon resonance (**Fig. 2E**) gave an apparent Kd of 137 +/-89 nM for Langya G and 41 +/-15 nM for Wenzhou G. The ∼3-fold difference between Langya and Wenzhou is almost entirely due to slower dissociation rather than increased association. As the G constructs are tetrameric, these are apparent, avidity-influenced constants. The increased binding affinity of Langya and Wenzhou relative to Mojiang is consistent with the reduced pseudovirus entry of Mojiang (**Fig 2A-C**).

Next, we tested whether protein C and a monoclonal antibody against EPCR (clone RCR- 401) were sufficient to block Langya, Wenzhou, and Mojiang entry. VSVΔG-luc coated with human EPCR was pre-incubated with 20 μg/mL recombinant human protein C, 10 μg/mL anti-EPCR, or PBS, and then added to 293T cells expressing the G and F proteins of each virus (**Fig 2F**). Protein C modestly, but statistically significantly, reduced Langya and Wenzhou G/F-mediated entry while it had a greater effect on Mojiang entry. Anti-EPCR more potently restricted entry of all three viruses. This suggests that Langya, Wenzhou, and Mojiang have at least a partially overlapping epitope with protein C and anti-EPCR clone RCR-401.

Our findings that EPCR is sufficient for Langya virus entry enabled us to develop a neutralization assay. First, we tested plasma from three rhesus macaques vaccinated against Langya G and Nipah G. Plasma from all three macaques neutralized both Langya and Wenzhou (**Fig. 2G**). While neutralization of Langya by homotypic NHP plasma was 4-fold more potent than that of Wenzhou (p<0.05, paired t-test), this demonstrates cross-reactivity within the Langya virus clade. Two previous monoclonal antibodies have been described against Langya (1069 and 1133)(*35*). However, it has previously not been possible to test neutralizing activity of these antibodies. Here we confirm that both antibodies bound Langya G expressed on the cell surface by flow cytometry. However, neutralizing activity was modest for both 1069 and 1133 (**Fig. 2H**). This highlights the need for development of future potent neutralizing mAbs against the Langya virus clade.

### Cryo-EM structure reveals mechanism of Langya G and EPCR interaction

To determine the structural basis of Langya G interactions with human EPCR, we resolved a 3.1 Å cryo-EM structure (**Figure S1-2 and Table S1**). This structure revealed a one-to-one binding stoichiometry of EPCR with each of the four Langya G head domains (**Fig 3A**). A density corresponding to the endogenous diacyl phospholipid phosphatidylethanolamine (PTY) was resolved in the MHC-like groove of EPCR. The Langya G-EPCR binding stoichiometry is the same as that for Nipah G and ephrin-B2(*36*); however, the receptor binding site and binding mode on Langya G is distinct and non-overlapping from that of Nipah G (**Fig 3B**). The antibody epitope of anti-Langya mAbs 1069 and 1133 are also remote from the EPCR binding interface consistent with limited neutralizing activity (**Fig. 3C and 2H**).

**Fig 3.**
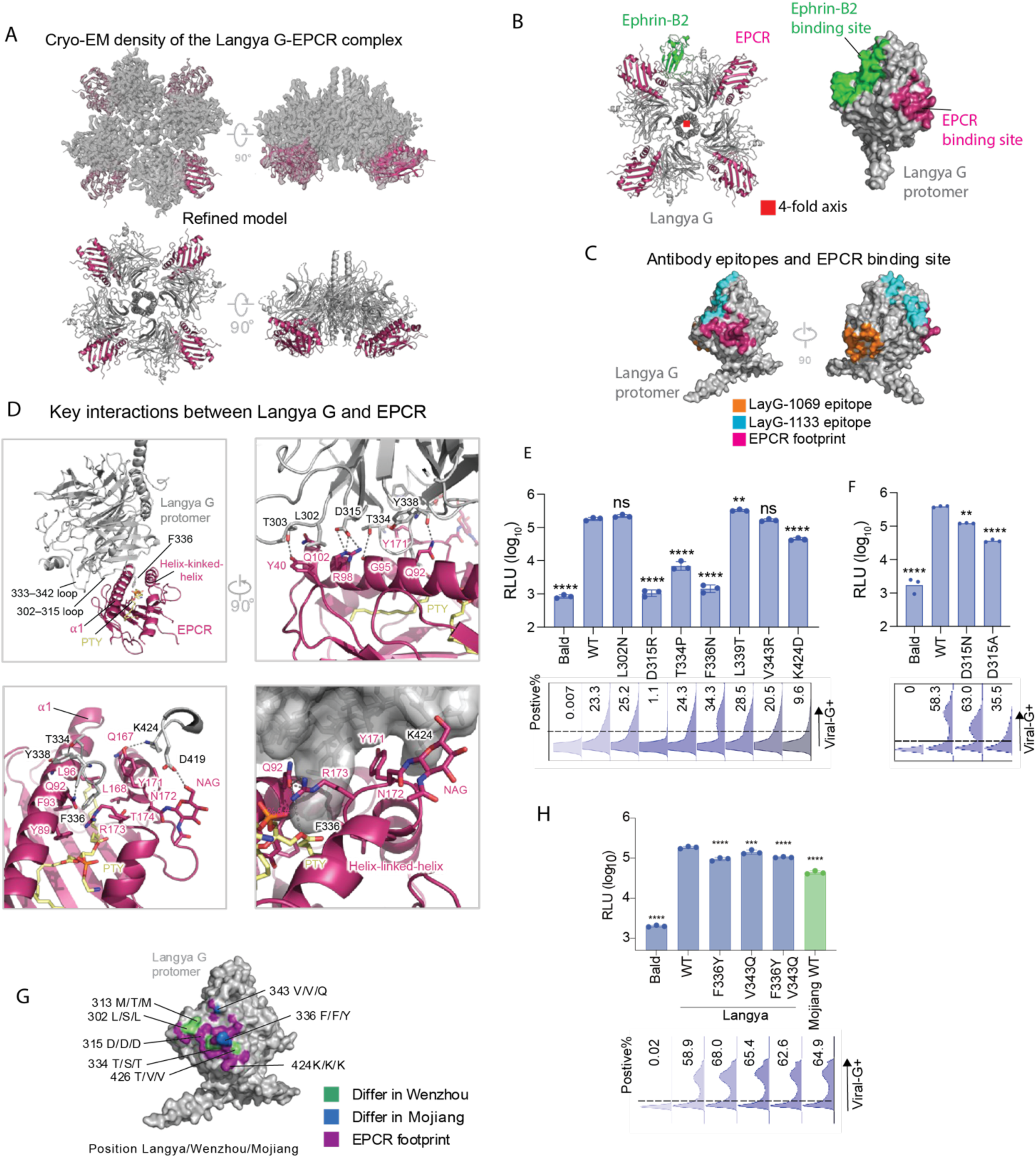
Cryo-EM structure reveals mechanism of Langya G and EPCR interaction. (**A**) 3.1 Å cryo-EM structure reveals EPCR (magenta) binds to the head domain on Langya G (gray) with 1:1 stoichiometry. (**B**) Left, the Langya G tetramer (gray) in complex with EPCR (magenta), viewed down the molecular four-fold axis (red square), with ephrin-B2 (green) from the Nipah G–ephrin-B2 complex (PDB 2VSM) superposed onto one Langya G head domain. Right, surface representation of a Langya G protomer, colored by the EPCR footprint (magenta) and by the residues structurally equivalent to the Nipah G ephrin-B2 site (green). The two receptor binding sites are non-overlapping. (**C**) EPCR footprint (magenta) on Langya G relative to the epitopes of the previously reported Langya G antibodies LayG-1069 (orange) and LayG-1133 (cyan), in two orientations related by a 90° rotation. (**D**) Key interactions of Langya G and EPCR are shown. (**E-**F) Single amino acid substitutions on Langya G residues identified at the EPCR interface were introduced. Each Langya G variant and WT Langya F were expressed in 293T cells and infected with VSVΔG-luc coated with human EPCR. Cell surface expression of each Langya G was assessed by flow cytometry with anti-flag. Entry was quantified by luciferase activity (RLU, relative light units). Bald refers to control without EPCR. (**G**) EPCR-contact residues mapped onto the surface of a Langya G protomer. The EPCR footprint is magenta and surface outside the footprint is gray. Within the footprint, positions at which the residue differs in Wenzhou G are green and those that differ in Mojiang G are blue; positions identical in all three viruses remain magenta. Labels give the residue number in full-length Langya G followed by the residue in Langya/Wenzhou/Mojiang. (**H**) Reciprocal substitutions between Langya G and Mojiang G were introduced, expressed on 293T cells along with cognate WT F, and then challenged with VSVΔG-luc coated with human EPCR. Data represents the mean of three independent experiments each performed in triplicate. Data is analyzed by one-way ANOVA with Dunnett’s multiple comparisons test comparing each column to WT Langya G control. Error bars denote mean+/-SD. *p<0.05, **p<0.01, *** p< 0.001, ****p<0.0001 Ns, not significant. PTY, diacyl phospholipid phosphatidylethanolamine.

The Langya G protein engages human EPCR by inserting the protruding F336 into the hydrophobic groove between EPCR α1 helix and the helix-kinked-helix. EPCR’s α1 helix docks into a cavity bounded by the I333–T343 and L302–D315 loops of Langya G, while the helix-kinked-helix is wedged into the cleft between F336 and K424 (**Fig. 3D. upper left**). The two proteins bury a comparable amount of surface area (BSA) on each side, roughly 858 Å² on EPCR and 884 Å² on Langya G, with smaller fractions attributable to an EPCR-linked glycan on N172 (NAG, 33 Å²) and an EPCR-bound phosphatidyl lipid (PTY, 48 Å²) in the lipid-loaded groove (**Table S2**).

The interface is stabilized by extensive hydrogen bonds and salt bridges. On the EPCR α1 helix, residues Y40, Q92, G95, R98, Q102 are key hydrogen-bonding partners which pair with Langya G residues T303, Y338, T334, F314 and R332, respectively. EPCR R98 is particularly important, not only forming a hydrogen bond to the backbone oxygen of F314 and salt bridges with D315 of Langya G, but also providing 10% of the BSA, making it one of the most central residues at the interface (**Fig. 3D, upper right, and Table S2**). On the helix-kinked-helix side, Q167 and the glycan residue NAG201 form hydrogen bonds with K424 and D419, respectively. EPCR Y171 stacks into the cleft between Langya G F336 and K424 and provides ∼14% of the BSA (**Fig. 3D, bottom and Table S2**).

The EPCR-binding surface on Langya G is centrally anchored on F336, which provides ∼24% of the BSA. The F336-binding pocket on EPCR is formed by hydrophobic residues Y89, F93, L96 and L168 from the α1 helix and the helix-kinked-helix. With the endogenous PTY lining at the bottom, the rim of the pocket is primarily patched by Q92, Q167, Y171 and R173, with R173 latched in place by hydrogen bonds from O4 atom of PTY and Q92 (**Fig. 3D, bottom and Table S2**). Together, Langya G binds EPCR via a compact, hydrogen-bond- and salt-bridge-stabilized interface built around the EPCR hydrophobic groove, with F336 acting as a central anchoring residue.

To validate the cryo-EM structure, we introduced substitutions in Langya G at seven sites (L302, D315, T334, F336, L339, V343, and K424) with high BSA in the EPCR complex (**Fig. 3E and Table S2**). We substituted each Langya G residue for the cognate residue in the non-pathogenic Cedar henipavirus which uses ephrins for entry(*37*, *38*), the only exception being position 315 which was Asp in both Langya and Cedar G and thus we substituted it initially with an Arg instead. We tested the effects of these substitutions in a reverse pseudotype assay with human EPCR. Cell surface expression of each Langya G variant was assessed by flow cytometry with anti-flag. Langya D315R failed to express on the cell surface, so we introduced D315N and D315A, both of which expressed. Langya G D315N, D315A, T334P, F336N, and K424D significantly reduced EPCR-mediated entry (**Fig. 3E-F**), consistent with the structural roles of these residues at the binding interface. Substitutions at D315 and T334 abolish salt bridges and hydrogen bonds with EPCR R98 and G95. K424D removes the hydrogen bond with EPCR Q167, while replacing F336 with the polar Asn disrupts interactions with the EPCR hydrophobic pocket. These results confirm that the hydrogen bonds, salt bridges, and the F336 anchor point identified in the cryo-EM structure are functionally required for EPCR-mediated entry.

To investigate the molecular basis for reduced binding and entry of Mojiang relative to Langya and Wenzhou, we evaluated 22 Langya G residues at the receptor binding interface (**Fig. 3G**). All 22 residues are perfectly conserved across the 11 unique Langya virus genomes(*3*, *17*).

Only two of the 22 residues are conserved between Langya G and Wenzhou G (V343 and F336) but not in Mojiang G (Q343 and Y336) (**Fig. 3F-G and Figure S3**). We introduced the Mojiang residues at these two sites alone and in combination in Langya G (F336Y and V343Q and tested infectivity by reverse pseudovirus infection. Substitution of Langya F336 with Tyr from Mojiang reduces infection by 2-fold with no additional synergistic effect in the F336Y/V343Q double substitution (**Fig 3H**). Consistent with this, structural modeling of the V343Q substitution in the Langya G-EPCR complex did not identify significant steric clashes. However, the F336Y substitution introduces a hydroxyl group that potentially causes steric clashes with pocket-lining residues, particularly L96 and PTY.

### EPCR molecular determinants of Langya, Wenzhou, and Mojiang virus entry

Next, we evaluated the EPCR determinants of parahenipavirus entry. The cryo-EM structure reveals 22 EPCR residues that contact Langya G with 17 of these residues having a buried surface area ≥10 Å² (**Table 2**). These contact residues have significant overlap with the binding sites of both protein C and PfEMP1 (**Fig 4A**). Interestingly, both human Protein C and PfEMP1 have a Phe residue (F4 in protein C and F656 in PfEMP1) that inserts into the same EPCR binding pocket as Langya G F336 (**Fig 4B**)

**Fig 4.**
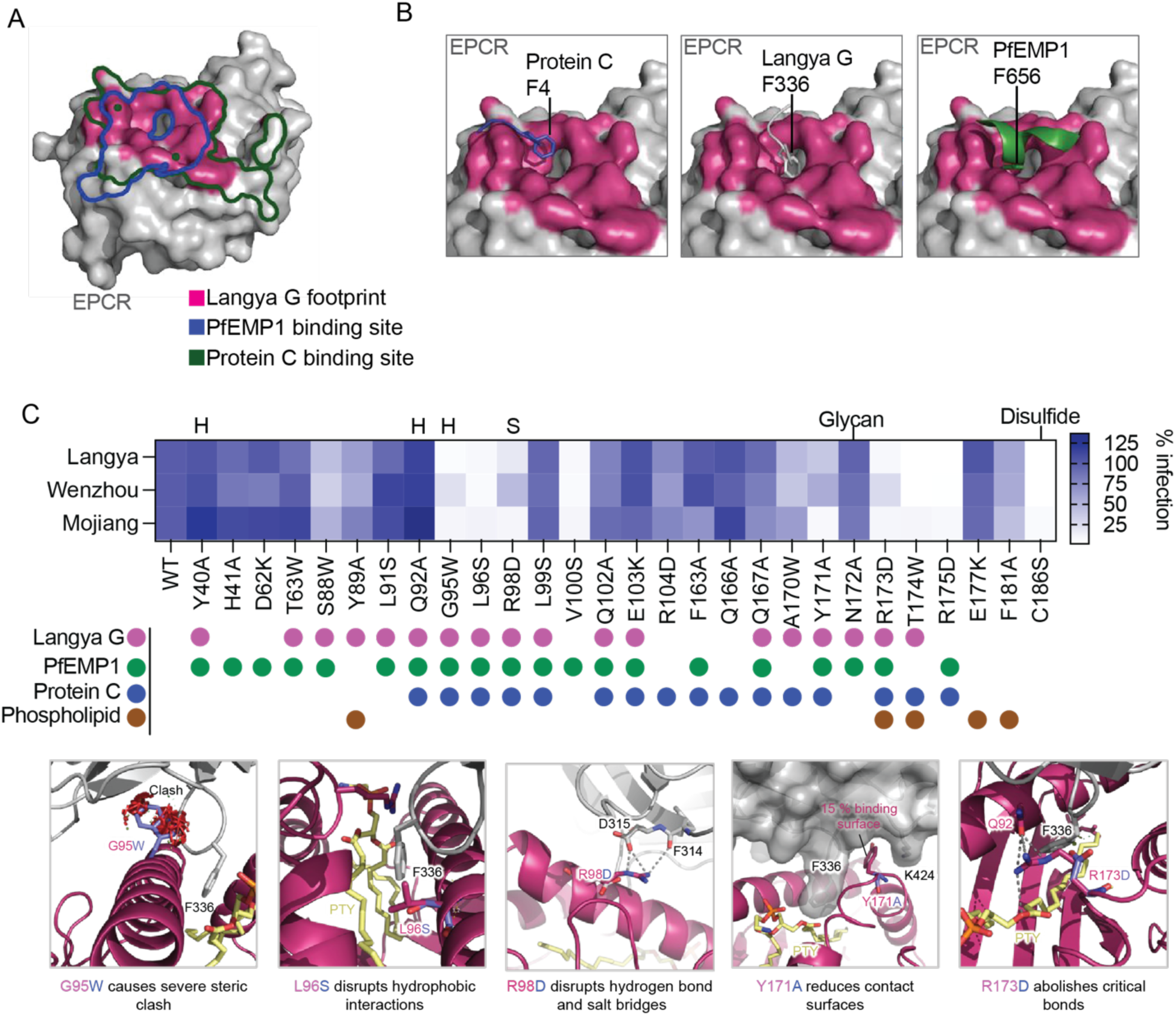
EPCR molecular determinants of Langya, Wenzhou, and Mojiang virus entry. (**A**) Surface representation of the human EPCR ectodomain showing the Langya G footprint (magenta) and that of protein C (blue) and PfEMP1 (green). (**B**) The respective Phe residues from protein C, Langya G, and PfEMP1 which interact with the same EPCR binding pocket are shown. (**C, top**) Entry of VSVΔG-luc coated with human EPCR (WT or point mutant) into 293T cells expressing the WT F and G proteins of Langya, Wenzhou or Mojiang virus, shown as a percentage of entry relative to WT EPCR. Each cell is the mean of four independent experiments. Annotations above the map indicate EPCR residues that form a salt bridge (S) and/or a hydrogen bond (H) with Langya G, a putative N-linked glycosylation site (Glycan) and the cysteine of a disulfide bond (Disulfide). Assignment of each residue to the Langya G footprint (magenta), the PfEMP1 footprint (green), the protein C footprint (blue) and the phospholipid-binding groove (brown) is shown below. (**C, bottom**) Key EPCR contact residues validated in (**C**, **top**) (G95, L96, R98, Y171, and R173) shown in the Langya G–EPCR complex together with their contacting residues in Langya G. PTY, diacyl phospholipid phosphatidylethanolamine.

**Table 2.** Interactions between EPCR and Langya G glycoprotein.

| Interface areas |  |  |  |  |  |  |  |
|---|---|---|---|---|---|---|---|
|  | Langya_G binding area on EPCR (Å²) | EPCR binding area on Langya_G (Å²) |  |  |  |  |  |
| Protein | 778.0 | 808.3 |  |  |  |  |  |
| EPCR glycan | 32.8 | 31.2 |  |  |  |  |  |
| EPCR PTY | 47.6 | 44.2 |  |  |  |  |  |
| Total | 858.4 | 883.7 |  |  |  |  |  |
| Hydrogen bonds and salt bridges between EPCR and Langya_G |  |  |  |  |  |  |  |
| Hydrogen bonds |  |  |  | Salt bridges |  |  |  |
| ## | EPCR | Dist. [Å] | Langya_G | ## | EPCR | Dist. [Å] | Langya_G |
| 1 | TYR40[OH] | 2.7 | THR303[OG1] | 1 | ARG98[NE] | 2.4 | ASP315[OD1] |
| 2 | GLN92[NE2] | 3.1 | TYR338[O] | 2 | ARG98[NH2] | 3.2 | ASP315[OD1] |
| 3 | GLY95[O] | 3.9 | THR334[OG1] |  |  |  |  |
| 4 | ARG98[NH2] | 3.3 | PHE314[O] |  |  |  |  |
| 5 | GLN102[OE1] | 2.4 | ARG332[NH1] |  |  |  |  |
| 6 | GLN167[OE1] | 3.8 | LYS424[NZ] |  |  |  |  |
| 7 | NAG201[O6] | 2.9 | ASP419[OD2] |  |  |  |  |
| Interface residues and buried surface areas |  |  |  |  |  |  |  |
| EPCR residues | Bonds | ASA (Å²) | BSA (Å²) | Langya_G residues | Bonds | ASA (Å²) | BSA (Å²) |
| TYR40 | H | 141.0 | 81.5\|\|\|\|\|\| | LEU302 |  | 58.0 | 51.0\|\|\|\|\|\|\|\|\| |
| SER84 |  | 68.6 | 7.9\|\| | THR303 | H | 75.9 | 26.1\|\|\|\| |
| GLN87 |  | 64.4 | 10.2\|\| | ALA304 |  | 51.2 | 9.2\|\| |
| SER88 |  | 50.4 | 19.4\|\|\|\| | ALA305 |  | 105.2 | 0.4\| |
| TYR89 |  | 63.8 | 10.5\|\| | MET313 |  | 96.5 | 37.1\|\|\|\| |
| LEU91 |  | 123.8 | 52.6\|\|\|\|\| | PHE314 | H | 31.2 | 17.0\|\|\|\|\|\| |
| GLN92 | H | 91.3 | 84.8\|\|\|\|\|\|\|\|\|\| | ASP315 | S | 41.4 | 33.5\|\|\|\|\|\|\|\|\| |
| PHE93 |  | 27.5 | 7.5\|\|\| | TRP318 |  | 68.1 | 10.8\|\| |
| HIS94 |  | 22.1 | 5.1\|\|\| | ARG332 | H | 95.0 | 55.7\|\|\|\|\|\| |
| GLY95 | H | 27.5 | 27.0\|\|\|\|\|\|\|\|\|\| | THR334 | H | 50.1 | 46.3\|\|\|\|\|\|\|\|\|\| |
| LEU96 |  | 52.5 | 34.5\|\|\|\|\|\|\| | GLY335 |  | 58.7 | 58.7\|\|\|\|\|\|\|\|\|\| |
| ARG98 | HS | 98.2 | 86.6\|\|\|\|\|\|\|\|\| | PHE336 |  | 216.0 | 216.0\|\|\|\|\|\|\|\|\|\| |
| LEU99 |  | 63.1 | 53.6\|\|\|\|\|\|\|\|\| | ALA337 |  | 21.5 | 18.9\|\|\|\|\|\|\|\|\| |
| GLN102 | H | 144.1 | 47.8\|\|\|\| | TYR338 | H | 58.0 | 49.3\|\|\|\|\|\|\|\|\| |
| GLN166 |  | 115.5 | 15.1\|\| | LEU339 |  | 120.0 | 37.7\|\|\|\| |
| GLN167 | H | 28.3 | 15.3\|\|\|\|\|\| | LEU340 |  | 29.8 | 4.4\|\| |
| LEU168 |  | 15.6 | 2.8\|\| | VAL343 |  | 86.1 | 35.5\|\|\|\|\| |
| ALA170 |  | 64.4 | 34.9\|\|\|\|\|\| | TYR344 |  | 12.9 | 7.6\|\|\|\|\|\| |
| TYR171 |  | 129.2 | 119.0\|\|\|\|\|\|\|\|\|\| | ARG372 |  | 177.0 | 7.3\| |
| ASN172 |  | 115.9 | 5.0\| | ASP419 | H | 68.8 | 35.0\|\|\|\|\|\| |
| ARG173 |  | 92.6 | 33.8\|\|\|\| | LYS424 | H | 160.9 | 102.0\|\|\|\|\|\|\| |
| THR174 |  | 38.8 | 23.4\|\|\|\|\|\|\| | PRO425 |  | 7.7 | 5.3\|\|\|\|\|\|\| |
| NAG201 | H | 356.1 | 32.8\| | THR426 |  | 50.6 | 18.9\|\|\|\| |
| PTY205 |  | 1238.3 | 47.6\| |  |  |  |  |

\*H and S: Residues making **H**ydrogen bond or **S**alt bridge; ASA: Accessible Surface Area, Å²; BSA: Buried Surface Area, Å²; \| Buried area percentage, one bar per 10%.

We performed targeted, structure-guided mutagenesis with the goal to induce maximally disruptive changes (i.e. charge reversals and steric clashes) at predicted EPCR interaction interfaces to ablate Langya entry. In addition to key Langya G contact sites, we also made EPCR substitutions at PfEMP1 binding sites outside of the Langya G binding surface (H41A, D62K, V100S, F163A, and R175D). We also mutated an N-linked glycan motif (N172A) near the Langya G binding site, five residues which support phospholipid binding (Y89A, R173D, T174W, E177, and F181A) and C186 which mediates an intramolecular disulfide bond with C118. We tested the ability of each EPCR mutant to support entry of Langya, Wenzhou, and Mojiang (**Fig 4C and Figure S4**). Individual substitutions predicted to disrupt the N-linked glycan (N172A) and phospholipid binding pocket outside of the Langya G footprint (E177K and F181A) had limited effect on viral entry, suggesting that the phospholipid-binding groove itself is dispensable for entry.

However, we observed two clusters of EPCR substitutions that abrogated entry of all three parahenipaviruses. The first cluster includes EPCR G95W, L96S, and R98D. G95 forms a hydrogen bond with Langya G T334 (**Fig 4C**). The G95W substitution causes severe clashes at the space constrained interface. As discussed earlier, R98D abolishes salt-bridges with Langya G D315 and hydrogen bond with F314, which were demonstrated to be critical for Langya G infectivity (**Fig 3E-F**). Modeling the R98D substitution onto the structure indicates that the charge reversal eliminates both interactions and brings D98 within 3.5 Å of D315 resulting in repulsion. The L96S change potentially reduced the hydrophobic interactions (**Fig 4C, bottom**). The second cluster includes Y171A, R173D, T174W, and R175D (**Fig 4C**). Y171 has the largest BSA of all EPCR contact residues, it inserts into the cleft between Langya G F336 loop and K424, providing ∼14% binding surface (**Fig 4C and Table S2**). R173, which was reported to act as a highly flexible conformational gatekeeper within EPCR, interacts directly with bound phospholipid PTY and Q92 to stabilize the hydrophobic pocket(*27*, *28*). Disrupting R173 interactions potentially affects the docking of Langya G F336 (**Fig 4C**). T174W introduces steric hindrance that impedes access to the EPCR pocket. Similarly, R175D would disrupt the kink region in the helix-kinked-helix.

We investigated space constraints around EPCR G95 at the Langya G-EPCR interface to determine which amino acids could be tolerated at this position without clashing with the Langya G protein (**Fig. S5**). Starting from the observation that a modeled Ser fit at this position, we systematically tested increasingly bulkier side chains to map the size and shape limits of the local binding interface pocket. Ser Cβ was at a tight but acceptable 3.6 Å from the G protein T334 Oγ atom, Thr was excluded because its β-branched substituent clashed severely (∼2 Å), indicating the interface couldn’t accommodate two heavy atoms at the Cβ position. Leu was tested to determine whether extending the branch point one bond further out (to Cγ) would provide sufficient rotational freedom — but it failed more severely still, with Cδ atoms overlapping the Langya G protein volume entirely, demonstrating that the interface constraint is governed by overall reach and bulk rather than branch position alone. This systematic approach revealed space constraints at the viral-receptor interface, suggesting that even seemingly conservative substitutions away from Gly, Ala or Ser would likely disrupt the precise geometry required for Langya G-protein engagement with EPCR.

C186S, which lies outside of the Langya G binding interface, also blocked viral entry, likely due to disruption of a critical disulfide bond in the MHC-like fold rather than direct effects on binding of Langya G. V100S was another substitution which reduced viral entry that was not directly at the binding interface; however, V100 is in the hydrophobic pocket and directly adjacent to key residues at the binding interface. The fact that Langya, Wenzhou and Mojiang were similarly affected across this EPCR substitution panel indicates that the three viruses engage an overlapping surface on EPCR rather than distinct sites on a shared receptor.

### EPCR sequence variation across animal species predicts parahenipavirus host range

Next, we asked whether natural variation at these EPCR positions govern EPCR host range. We assembled EPCR orthologs from 26 vertebrate species including natural reservoirs of parahenipaviruses including four shrew, five rodent, five bat EPCRs along with human adjacent animals that could represent intermediate hosts such as livestock. Outlier species are also included to maximize EPCR genetic diversity. We used a reverse pseudotype assay in which we expressed Langya, Wenzhou, or Mojiang G/F on the surface of 293T cells and then challenged cells with VSVΔG-luc coated with each animal EPCR or a no EPCR (bald) control (**Fig. 5A**). Orthologs from all four shrew species (S*uncus etruscus, Sorex araneus, Crocidura gueldenstaedtii* and *Blarina brevicauda*), from the star-nosed mole Condylura cristata, and from human supported the highest entry of all three viruses. Orthologs from house mouse, cat, rhesus macaque, golden hamster, the big brown bat Eptesicus fuscus, bank vole and wood mouse supported intermediate entry. The remaining thirteen orthologs including four of the five bats, all three ungulates, dog, Norway rat, elephant, hedgehog, opossum and chicken were at or near background. There was strong concordance in EPCR use across the Langya, Wenzhou, and Mojiang with Mojiang entry being broadly less efficient, consistent with that in Fig 2.

**Fig 5.**
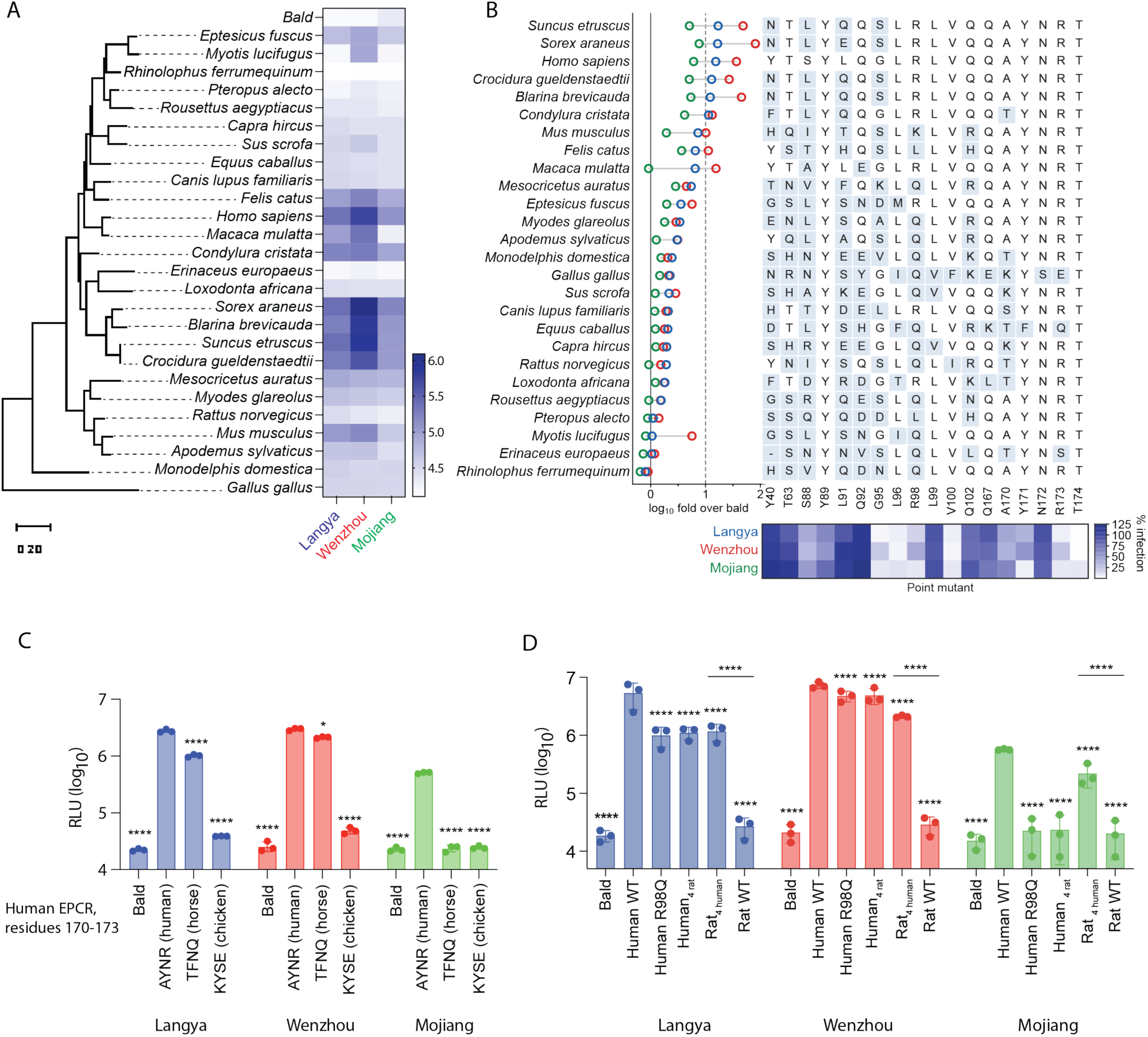
EPCR sequence variation across animal species predicts parahenipavirus host range. (**A**) Maximum-likelihood phylogeny of EPCR orthologs from 26 vertebrate species (left) and infectio of 293T cells expressing Langya, Wenzhou or Mojiang G and F by VSVΔG-luc coated with each EPCR ortholog. Bald particles lack EPCR. Heat map, log10 RLU. Scale bar, 0.2 substitutions per site. (**B**) Left, the data in panel (**A**) expressed as log10 fold over the no-EPCR control for Langya (blue), Wenzhou (red) and Mojiang (green); species are ordered by susceptibility. Middle, key EPCR residues that contact or promote Langya G entry, numbered according to full-length human EPCR. Residues that differ from human EPCR are shaded blue. Bottom, percent infection of human EPCR point mutants at these positions (from Fig. 4C) relative to WT human EPCR. (**C**) Infection of cells expressing Langya (blue), Wenzhou (red) or Mojiang (green) F and G by VSVΔG-luc coated with human EPCR bearing the native human EPCR 170-173 motif (AYNR) or that from horse (TFNQ) or chicken (KYSE). The leftmost bar in each group lacks EPCR. (**D**) Infection of 293T cells expressing Langya, Wenzhou or Mojiang G and F by VSVΔG-luc coated with EPCR. We include WT human, WT rat (*Rattus tanezumi*), human EPCR with a single rat residue (R98Q) or with four rat residues (human_4 rat_: H94N/G95S/R98Q/A170T). Four human residues were introduced into rat EPCR (rat_4 human_: N94H/S95G/Q98R/T170A). Data in (**A**) is the mean of two independent experiments, each performed in triplicate. Data in (**C**) were analyzed by one-way ANOVA with Dunnett’s multiple comparisons test against WT human EPCR within each virus. Data in (**D**) were analyzed using one-way ANOVA with Dunnett’s multiple comparisons test against WT human EPCR, with an additional comparison between rat WT and rat_4 human_ EPCR. *P < 0.05, ****P < 0.0001. RLU, relative light units. Bald, VSVΔG-luc without EPCR.

Interestingly, EPCR susceptibility did not track with host phylogeny. The European hedgehog is in the same eulipotyphla order as shrews and the star-nosed mole, yet hedgehog EPCR did not support entry. Conversely, EPCR from house mouse supported intermediate entry while that of the Norway rat was at background, and among the five bats tested only *Eptesicus fuscus* supported significant entry across all three viruses.

Instead, viral susceptibility tracked with sequence identity at the EPCR interface as a whole rather than at any single contact position. EPCR orthologs that support entry retain the human and shrew residues at key contact positions (Q92, G/S95, R98, and A170) demonstrated to be important for entry (**Fig. 4**), whereas non-susceptible EPCR orthologs are substituted at one or more of these sites (**Fig. 5B**).

Next, we tested EPCR interface residues between humans and non-susceptible animal EPCRs to reveal the molecular basis of species compatibility. First, we replaced the human EPCR residues 170-173 (AYNR) with the corresponding motif from horse (TFNQ) or chicken (KYSE) which are not susceptible to Langya, Wenzhou, or Mojiang (**Fig. 5C**). The chicken motif reduced entry of all three viruses to background levels. The horse motif modestly reduced Langya and Wenzhou entry, but reduced Mojiang to background. This parallels the critical role of Y171 observed in human EPCR (**Fig. 4C**). Thus, transfer of one to four residues from a non-permissive EPCR ortholog (i.e. chicken or horse) is sufficient to reduce receptor function.

Next, we compared human and rat EPCR which was not susceptible to Langya, Wenzhou, or Mojiang (**Fig. 5D**). We synthesized EPCR of the putative Mojiang reservoir *Rattus tanezumi* rather than that of *Rattus norvegicus* used in Fig 4B(*39*). *Rattus norvegicus* and *Rattus tanezumi* EPCR are >98% identical and conserved at every contact position (except position 88; Ile in *R. norvegicus* and Leu in *R. tanezumi*) shown in (**Fig 4C**). We introduced reciprocal substitutions between human and rat EPCR at positions 94, 95, 98, and 170. Introduction of rat residues into human EPCR at position 98 alone or at all four positions in combination reduced infection of Langya, Wenzhou, and Mojiang (**Fig. 5D**). Similarly, introduction of the human residues at 94, 95, 98, and 170 into rat RPCR conferred susceptibility of rat EPCR to all three parahenipaviruses.

## Discussion

Here, we identify EPCR as an entry receptor for Langya, Wenzhou, and Mojiang viruses. This binding mechanism is distinct from the ephrin-mediated entry of Nipah and Hendra viruses indicative of substantial evolutionary change in receptor use within the virus family. Langya G engages a hydrophobic pocket also recognized by protein C and the malarial adhesin PfEMP1. These structurally and evolutionarily unrelated ligands converge on a common receptor surface through similar chemical interactions mediated by a key ligand Phe. The role of EPCR in protein C activation suggests that this surface is functionally constrained, potentially limiting host escape through receptor variation. Genetic loss of EPCR causes placental thrombosis and is embryonically lethal(*40*). Competition for this pocket also raises the possibility that viral engagement perturbs EPCR function and thus activated protein C(*29*).

Our findings strengthen the rationale for expanded surveillance of the Langya virus clade including Wenzhou and Mojiang viruses. Together with detection of Langya virus in wild shrews, the ortholog data support shrews as likely reservoir hosts. No EPCR sequence is available for the shrew species in which these viruses have been found (i.e. *Crocidura lasiura* and *Crocidura shantungensis* for Langya virus and *Suncus murinus* for Wenzhou virus), so the four shrew orthologs tested here, serve as proxies for the receptors of the natural hosts. Importantly, EPCR compatibility extends beyond recognized animal reservoirs informing cross-species transmission potential of these viruses into novel animal hosts. All 22 EPCR contact residues in Langya G were conserved across all 11 available viral genomes, including those from human and shrews from China and South Korea(*3*, *17*). The 35 human cases identified in the initial surveillance study likely substantially underestimate the burden of infection, because surveillance was restricted in time, geography, and clinical presentation(*3*). Detection in shrews beyond the areas of recognized human disease further supports broader surveillance and serosurveys of exposed populations.

Infectious Langya virus has not been successfully cultured, and an animal model of infection remains unavailable. Identifying EPCR provides a biological foundation for developing these systems, which are needed to investigate tissue tropism, disease pathogenesis, and evaluate countermeasure efficacy. Although multiple orthogonal assays establish the role of EPCR in glycoprotein-mediated entry, the necessity of EPCR during authentic infection and the possibility of alternative receptors remain unresolved. Notably, Daeryong, Melian, Ninorex, Gamak, Salt Gully, and Angavokely viruses did not use human EPCR, leaving the receptor use of these viruses unclear.

Mojiang virus used human EPCR least efficiently across both viral entry and binding assays. This informs the proposed association with the 2012 pneumonia deaths among miners(*5*). Viral RNA was recovered from rats at the mine months later, without direct evidence of infection in the patients. Inefficient receptor use does not exclude human infection but cautions against describing Mojiang virus as a definitive human pathogen. In addition, the inability of Mojiang to use rat EPCR from *Rattus norvegicus* and *Rattus tanezumi* suggests rats may not be the natural reservoir host. It is also possible that the deposited Mojiang G sequences have sequencing errors, that Mojiang uses a different receptor in its putative reservoir host (*Rattus tanezumi*), or that authentic Mojiang virus can tolerate reduced G binding affinity and fusogenicity. We also observed that Mojiang was more sensitive to mutations in both G and EPCR. This is consistent with lower affinity and infectivity relative to Langya and Wenzhou as may be easier to perturb the weak intermolecular interactions of Mojiang G and EPCR.

Importantly, this work enables improved countermeasure development and surveillance efforts against this important clade of viruses. Identification of EPCR enabled the development of neutralization assays which can distinguish antibody binding from functional inhibition, as illustrated by two previously described anti-Langya G antibodies that bound but poorly neutralized(*35*). Cross-neutralization of Wenzhou by Langya vaccine plasma suggests a single vaccine may confer protection within the Langya subclade. Despite the absence of reported human Wenzhou virus infections, the receptor compatibility and circulation in mammals living near people warrant active surveillance alongside Langya. Identifying EPCR enables countermeasure development through functional assays and informs surveillance efforts regarding susceptible hosts to mitigate zoonotic risk of this emerging virus clade.

---

**Figure S1.**
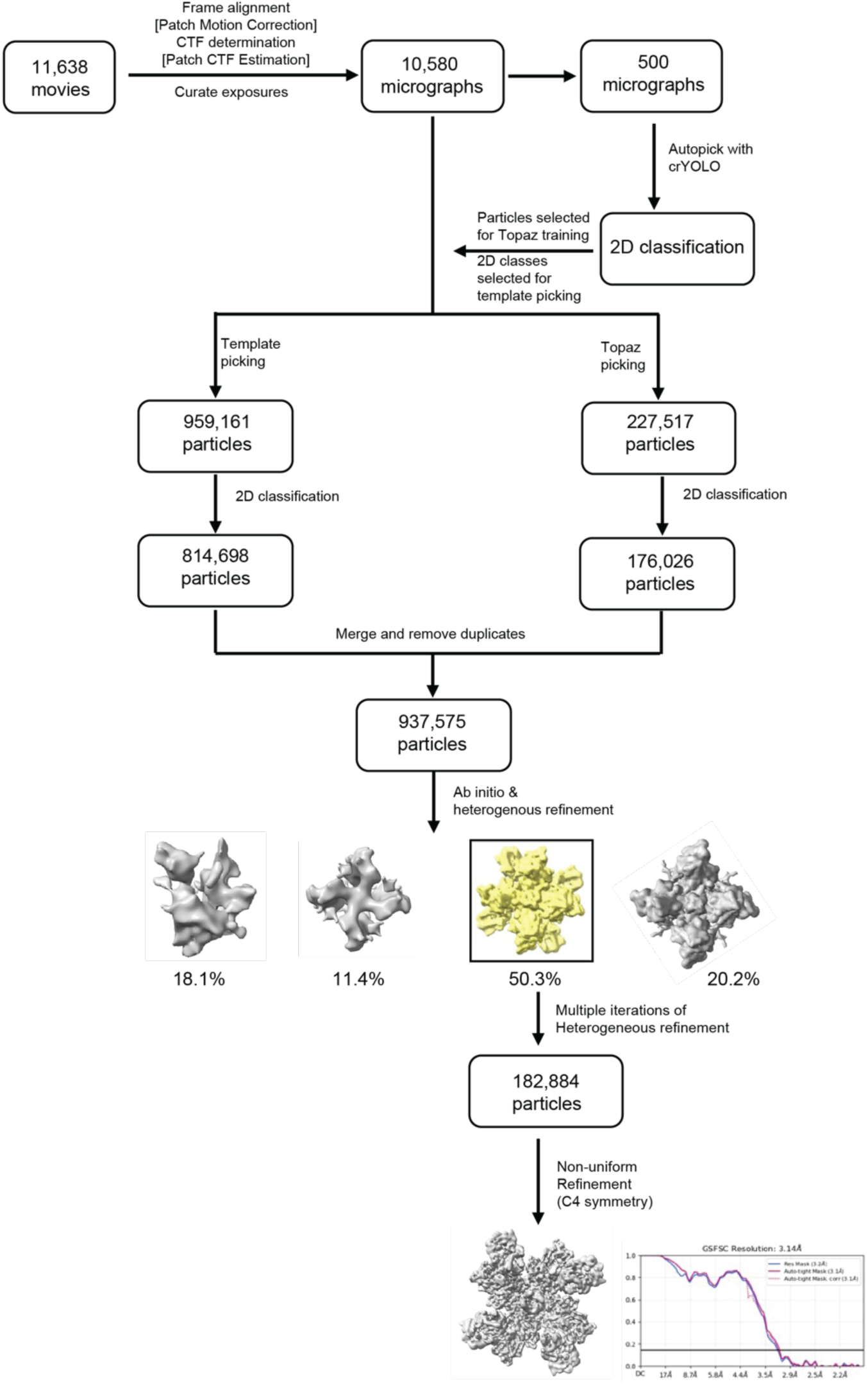
Cryo-EM data processing workflow for the Langya G – EPCR complex. Movies collected at 35° and 45° stage tilt were motion-corrected, CTF-estimated and curated in cryoSPARC. Particles picked from a 500-micrograph subset with crYOLO provided the 2D templates and the training set for Topaz; template-based and Topaz picks from the full dataset were merged and duplicates removed. Ab initio reconstruction followed by iterative heterogeneous refinement isolated the class boxed in yellow, and non-uniform refinement with C4 symmetry imposed gave the final reconstruction at 3.1 Å (gold-standard FSC = 0.143).

**Figure S2.**
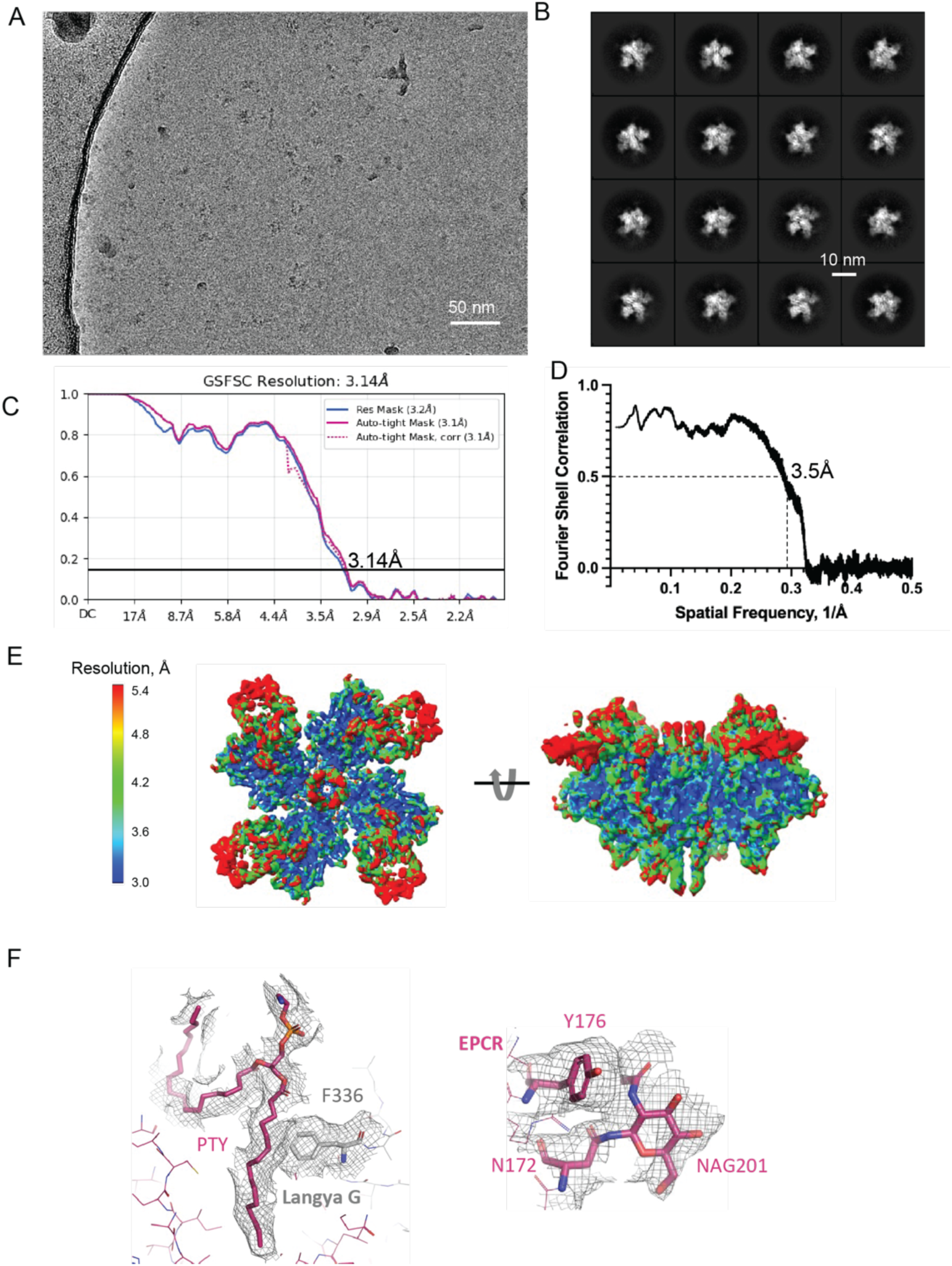
Validation of the cryo-EM reconstruction of the Langya G-EPCR complex. (**A**) Representative cryo-EM micrograph. (**B**) Representative high-resolution 2D class averages. (**C**) Gold-standard Fourier shell correlation (FSC) curve generated in cryoSPARC. The final resolution reached 3.1 Å at the FSC = 0.143 criterion. (**D**) FSC curve between the cryo-EM map and the refined atomic model. (**E**) Local resolution analysis calculated using ResMap(*41*). The cryo-EM map is colored according to local resolution. (**F**) Representative cryo-EM density at the interface between Langya G and EPCR.

**Figure S3.**
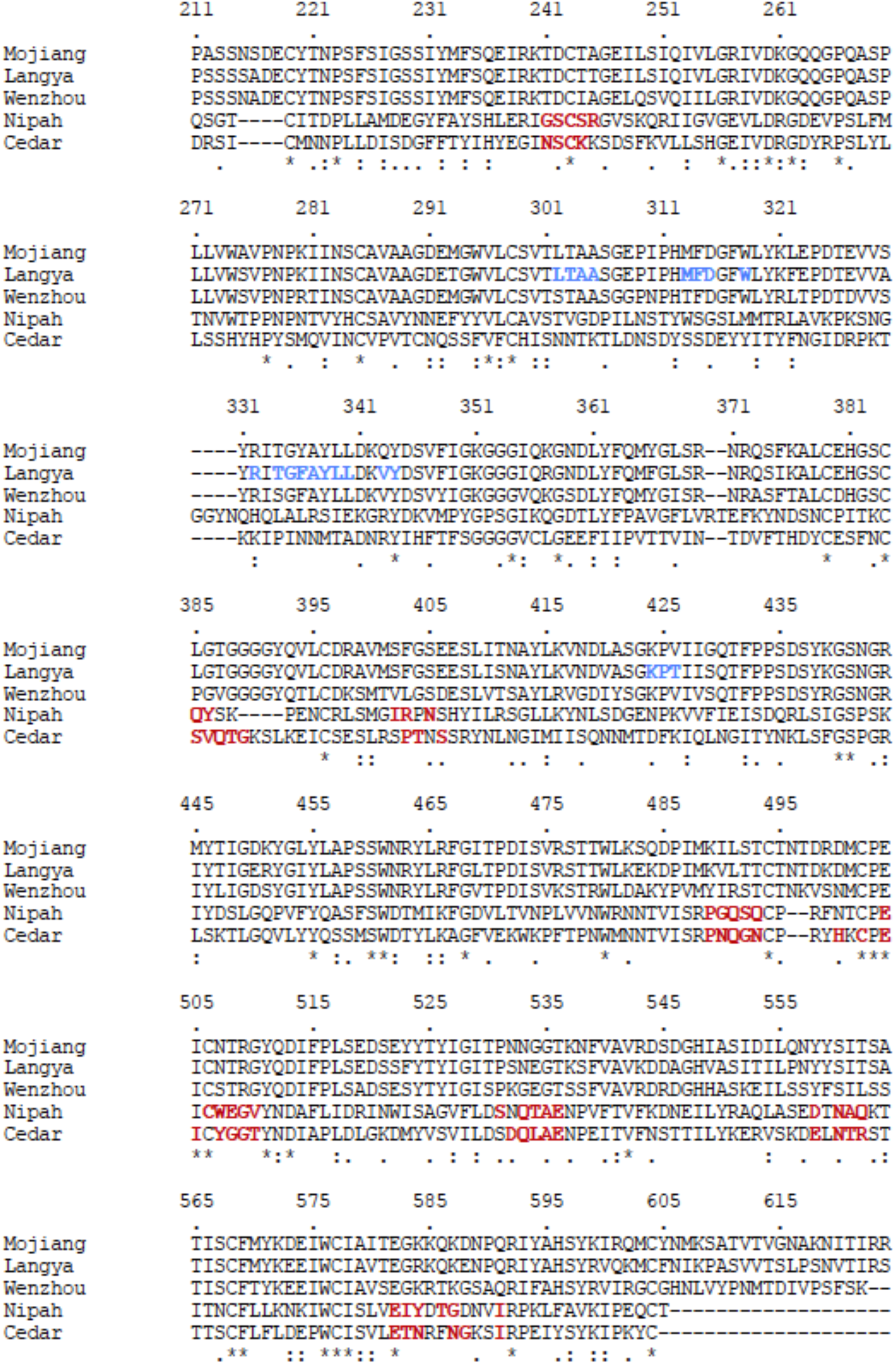
Alignment of Langya, Mojiang, Wenzhou, Nipah, and Cedar virus G head domains. Ephrin-B2 interacting residues for Nipah and Cedar G are highlighted in red and based on PDB 2VSM and 6P7Y, respectively. EPCR interacting residues on Langya G are highlighted in blue. Residue numbers corresponds to position in the full-length Langya virus G protein.

**Figure S4.**
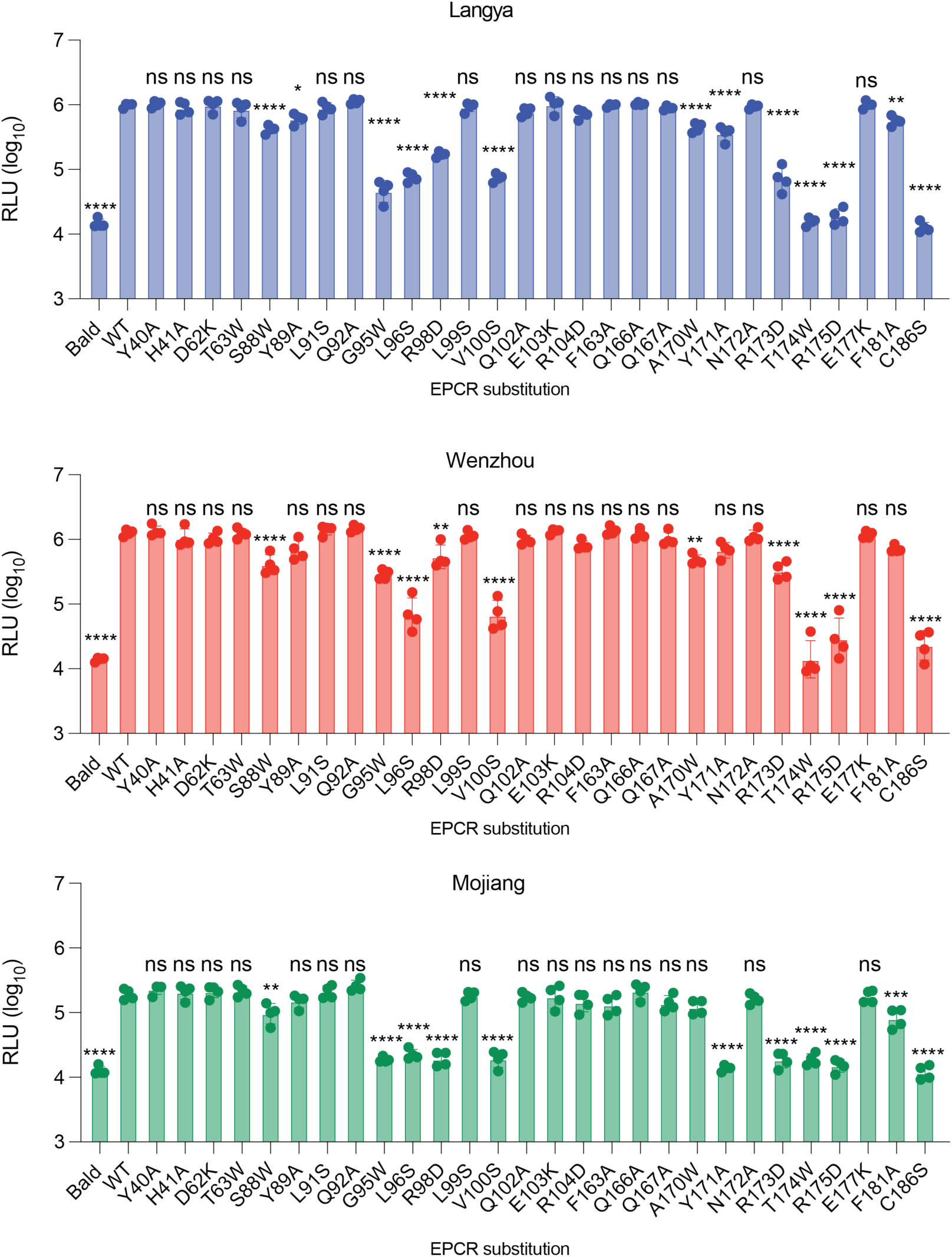
Unnormalized individual EPCR mutagenesis infection data with Langya, Wenzhou, and Mojiang related to Fig 4C. 293T cells were transfected with Langya G/F (top), Wenzhou G/F (middle), or Mojiang G/F (bottom) and challenged with VSVΔG-luc coated with human EPCR (WT or point mutant). Bald refers to VSVΔG-luc without EPCR. Infection was quantified by relative light units (RLUs). Data represent the mean of each of four independent experiments each performed in triplicate. Error bars, SD. Data analyzed by one-way ANOVA with Dunnett’s multiple comparison test. ****p<0.0001 ***p<0.001, **p<0.01, *p<0.05, ns, not significant.

**Figure S5.**
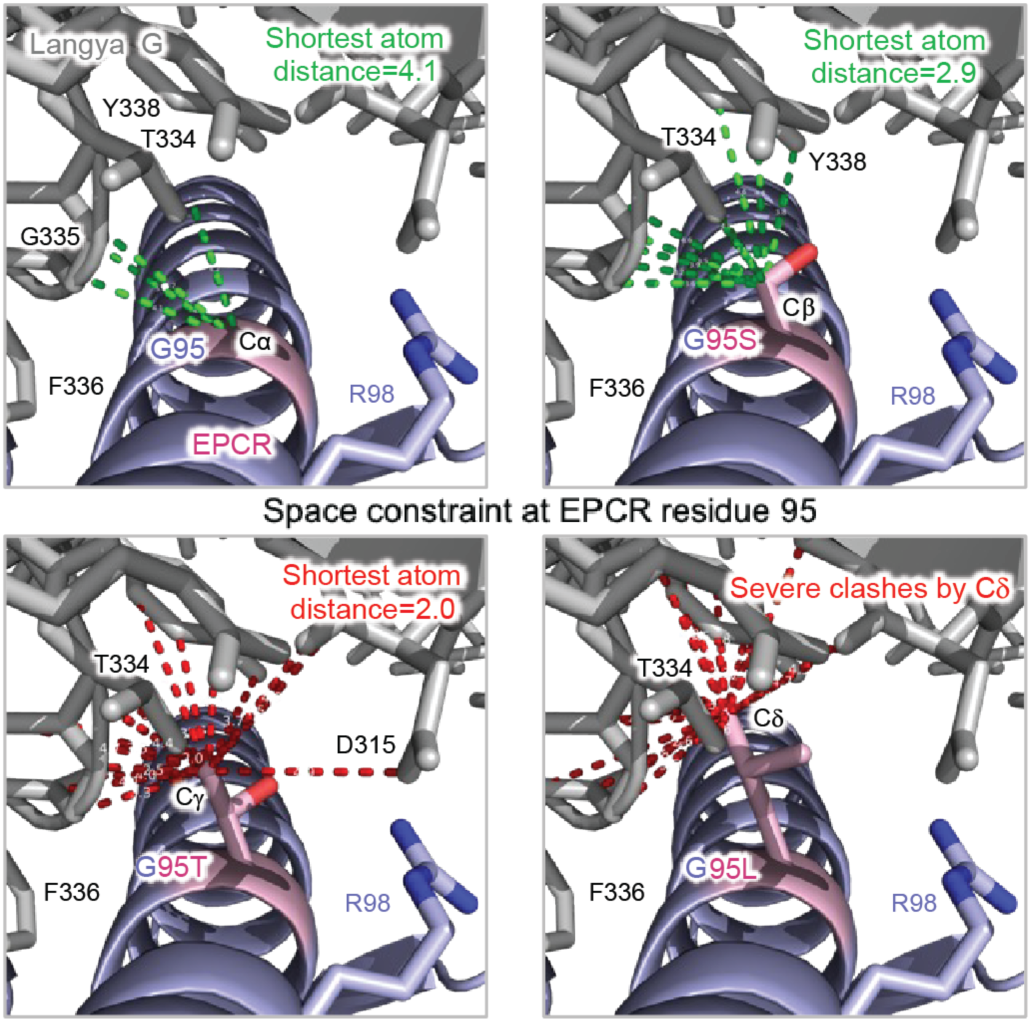
Structural interpretation of EPCR position 95. Molecular interactions of human EPCR G95 and Langya G. EPCR orthologs can tolerate a G or S at position 95 (top panels) but larger residues including Thr and Leu (bottom panels) introduce space constraints hindering G-EPCR interactions.

**Table S1.**
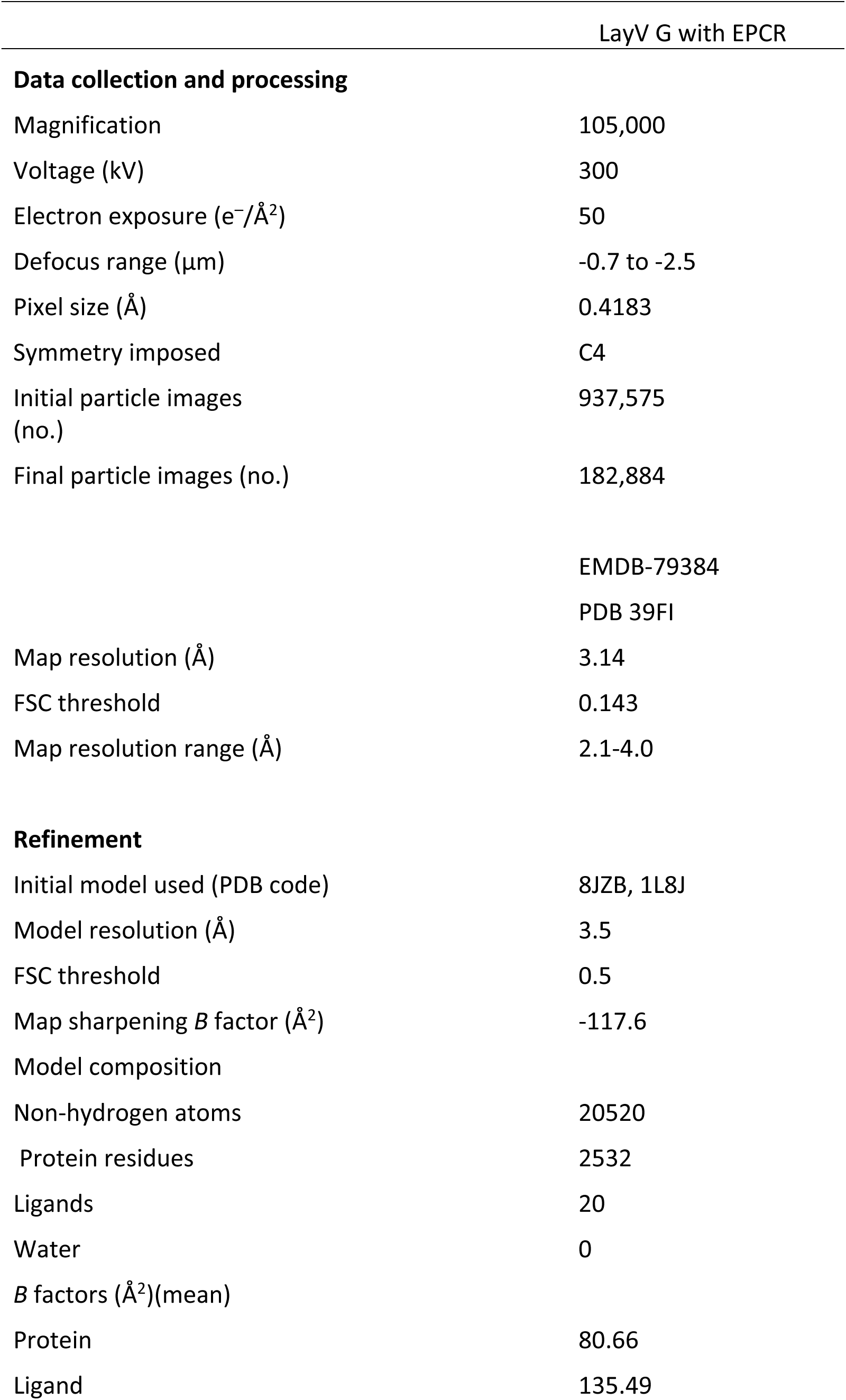
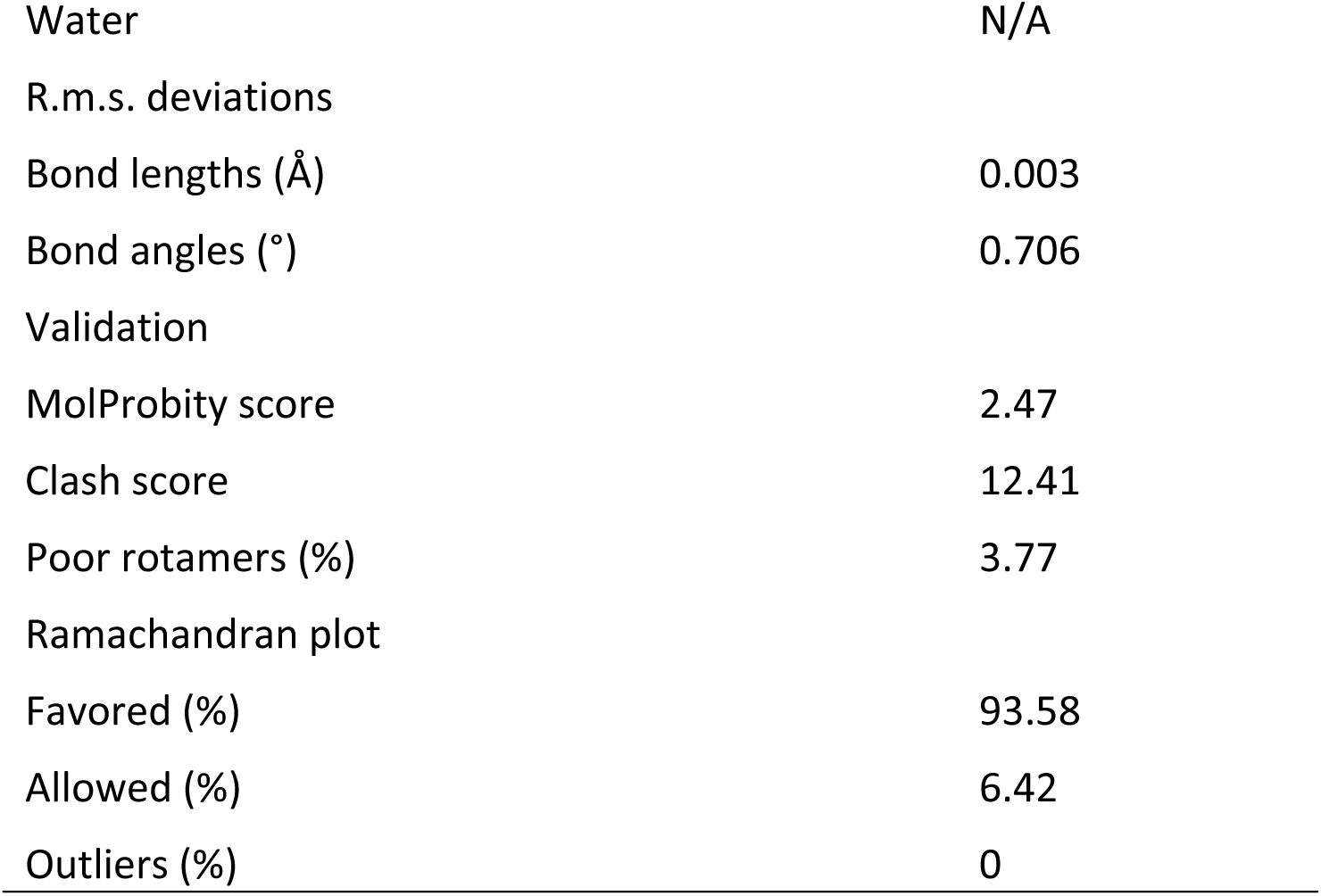
Cryo-EM refinement statistics.

**Data S1**. Metadata table of viruses with sequences of F and G proteins

**Data S2.** Human ephrin sequences

**Data S3.** CRISPR screen results

**Data S4.** EPCR ortholog sequences

## Methods

### Sequence and phylogenetic analysis

Protein sequence alignments were generated using MAFFT(*42*). For phylogenetic analyses, amino acid sequences of various EPCR or parahenipavirus attachment glycoprotein (G) sequences were aligned using MAFFT, and maximum-likelihood phylogenetic trees were inferred using MEGA12(*43*) with 500 bootstrap replicates. The amino acid substitution model and associated parameters were selected according to the best-fit protein model identified by MEGA12. The reservoir host, country of detection, accession number and G sequence for each virus is listed Supplementary Data 1.

### Plasmids

Human codon-optimized sequences encoding ephrins, EPCR orthologs, and EPCR mutants were synthesized with a C-terminal HA tag and subcloned into the pUC57 expression vector using BsmBI restriction sites(*34*).Human codon-optimized sequences encoding henipavirus and parahenipavirus attachment glycoproteins (G), G mutants, and fusion glycoproteins (F) were commercially synthesized and subcloned into the pCAGGS expression vector using EcoRI and NotI restriction sites, as previously described(*34*). G proteins contained a C-terminal FLAG tag.

### Fc-fusion protein production

A construct encoding the Langya virus attachment glycoprotein (G) head domain (residues 176–602) fused to a human IgG Fc domain (hFc-Langya-G) was commercially synthesized and produced by Twist Bioscience. Variable-region sequences of the LayG-1069 and LayG-1133 antibodies(*35*) were obtained from the structural model deposited in the Protein Data Bank (PDB: 9LEO) and used to generate recombinant human Fc-fusion proteins, which were commercially produced by Twist Bioscience.

### CRISPRa screen

The A375 CRISPRa surfaceome library was made as described in Song et al(*24*). Briefly, 5,891 genes were identified as encoding putative surface expressed proteins based on merging four datasets. Up to four guide RNAs were designed for each gene and 1000 non-targeting controls were included, cloned into the CRISPRa lentiviral plasmid pXPR_502 (Addgene, #96923), and transduced into A375 dcas9 VP64 cells(*44*).

A total of 32 million A375 CRISPRa surfaceome library cells were trypsinized, blocked with 2% FBS and 1mM Ethylenediaminetetraacetic acid (EDTA) in phosphate buffered saline (FACS buffer) for 45 min, incubated with hFc-Langya-G at 50 µg/ml for 45 min at 4C, washed in FACS buffer, incubated with rat anti-human IgG FITC secondary antibody (Biolegend #410720) for 30 min at 4C, and washed twice in FACS buffer. The positively stained cells were then live sorted through a 100 uM on the Sony MA900 Cell Sorter housed in a biosafety cabinet. A total of 14,437 FITC positive cells were sorted into a FACS tube and was cultured in a 96 well plate for 1 week before being expanded. DNA was isolated from the unsorted and sorted cells using the Zymo Genomic DNA Clean and Concentrator kit (Zymo) after proteinase k and RNase treatment. Isolated genomic DNA was then PCR amplified using standard staggered forward P5 primers and P7 reverse primers. Up to 10 ug of genomic DNA was mixed with 0.2 uM of forward primer mix and reverse primer with 50 ul 2x Q5 High-Fidelity master mix (NEB). PCR cycling conditions were 98 C 3 min, then 34 cycles of 98C 10s, 68C 20s, 72C 20s and final extension 72C 5 min. Gel extraction was performed to obtain pure PCR product using the Zymoclean Gel DNA Recovery kit (Zymo) and samples were sent with library preparation and sequencing to the Yale Center for Genome Analysis for sequencing on Illumina MiSeq with 150 bp paired end reads.

Reads were then matched and counted with the corresponding gene and guide number and the resulting count matrix was submitted to the Broad Apron analysis tool (https://portals.broadinstitute.org/gpp/public/analysis-tools/crispr-gene-scoring) for enrichment and depletion analysis (job 2026-10-03_09:05:39_059-broad). Settings were as follows: analysis type Hypergeometric, directionality Both, top n% guides per gene 100, dummy gene set size 4, min/max guides per gene 2/4, no-site controls displayed, 5 genes labelled. Non-targeting sgRNAs were renamed to the portal’s NO_SITE_ convention so they were recognized as no-site controls; Apron pooled the 1,000 of them into 249 four-guide control pseudo-genes. 6,138 genes were scored.

Argon P5 forward mix: px52_F1: 5’-ACACTCTTTCCCTACACGACGCTCTTCCGATCTCTTGTGGAAAGGACGAAACACCG - 3’ px52_F2: 5’-ACACTCTTTCCCTACACGACGCTCTTCCGATCTACTTGTGGAAAGGACGAAACACCG -3’ px52_F3: 5’-ACACTCTTTCCCTACACGACGCTCTTCCGATCTTCCTTGTGGAAAGGACGAAACACC G -3’ px52_F4: 5’-ACACTCTTTCCCTACACGACGCTCTTCCGATCTACTCTTGTGGAAAGGACGAAACAC CG -3’

Kermit P7 reverse primer kermit_R 5’

GTGACTGGAGTTCAGACGTGTGCTCTTCCGATCTTCTACTATTCTTTCCCCTGCACTG

T -3’

### Cell line production and culture

A375, HEK-293T, Hela-USU, and BHK-21 cells were maintained in Dulbecco’s modified Eagle medium (DMEM; Gibco) supplemented with 5% heat-inactivated fetal bovine serum (FBS) and 1% penicillin–streptomycin (Pen/Strep). Cells were maintained at 37 °C in a humidified atmosphere containing 5% CO₂.

To generate A375dCas9 EPCR CRISPRa cells, guides were ordered as oligos from Yale Keck Oligos service, annealed, phosphorylated, and cloned into pXPR_502(*45*): EPCR CRISPRa guide 1 CTCATTTCCTACTAGGGGGC and EPCR CRISPRa guide 2 GTGTCTCATTTCCTACTAGG and empty vector guide as control.

Lentivirus was produced by transfecting the plasmid with helper plasmids into HEK293T cells. Lentivirus was collected after 48-72 hours, clarified with centrifugation and passed through 0.45 uM filter. A375 dcas9 VP64 cells were then spinfected with lentivirus at 1,200g for 45 min. After 48 hours, cells were selected with puromycin 1 µg/ml. Cells were maintained in media with blasticidin 5 µg/ml and puromycin 1 µg/ml for all subsequent experiments.

To generate BHK cells stably expressing human EPCR, codon optimized human EPCR was cloned into pGS-LVV-puro by Genscript. Lentivirus was then produced as described above and transduced into BHK-21 cells with puromycin selection to generate EPCR-overexpressing BHK- 21 cells.

### Flow cytometry

Flow cytometry was used to assess parahenipavirus G expression, EPCR expression, reactivity of the LayG-1069 and LayG-1133 recombinant antibodies, and GFP positivity. For surface staining, transfected cells were collected and washed with flow cytometry staining buffer. Cell-surface G proteins were detected using the 5 µg/ml of LayG-1069 or LayG-1133 recombinant Fc proteins, or 5 µg/ml of anti-Flag tag antibody (Biolegend, #637302). Cells were subsequently stained with the appropriate fluorophore-conjugated secondary antibodies (1:300, v/v): FITC-conjugated anti-human IgG Fc antibody (BioLegend, #410720) for LayG-1069 and LayG-1133, or FITC-conjugated goat anti-rat IgG antibody (BioLegend, #405404) for the FLAG-tag antibody. Cells stained with the corresponding isotype or secondary-antibody-only controls were used to define background fluorescence. GFP expression was measured directly without antibody staining. Samples were acquired on a Beckman Coulter CytoFLEX S flow cytometer and analyzed using FlowJo v10.

### Production of VSV pseudoviruses

Vesicular stomatitis virus (VSV)-based pseudotyped viruses were generated as previously described(*34*, *46*), with modifications. Briefly, HEK-293T cells were transfected using polyethylenimine (PEI) with 5 µg each of pCAGGS plasmids encoding the G and F proteins from the same parahenipavirus for conventional (forward) pseudotyping, or with 10 µg of pUC57 plasmids encoding the indicated EPCR proteins for reverse pseudotyping. The transfection medium was replaced with fresh medium 5 h after transfection. At 24 h post-transfection, cells were inoculated with a replication-defective VSV vector in which the VSV-G open reading frame was replaced by a Renilla luciferase (luc) reporter cassette (VSVΔG-luc). After incubation for 1 h at 37 °C, the inoculum was removed, cells were washed twice with 1× PBS, and fresh culture medium was added. Pseudotyped virus-containing supernatants were collected at 24 and 48 h post-infection (hpi) and clarified by centrifugation to remove cellular debris. Clarified supernatants were concentrated tenfold using Amicon Ultra centrifugal filters with a 100-kDa molecular-weight cutoff (MWCO). Medium supplemented with the anti-VSV-G monoclonal antibody I4 (Kerafast) was subsequently added to neutralize residual input VSV-G-complemented virus, as previously described(*34*). Pseudotyped virus preparations were stored at −80 °C until use.

### VSV pseudotyped virus entry assays

For pseudotyped virus entry assays, 6 × 10⁵ HEK- 293T or BHK-21 cells were transfected with 2 µg of plasmid encoding the indicated EPCR ortholog or mutants, or 1 µg each of plasmids encoding the G and F proteins from the same parahenipavirus. Transfected cells were seeded at 6 × 10⁴ cells per well in 100 µl of culture medium in black-walled, clear-bottom 96-well plates (Corning #3904) for reverse pseudotyping assays and 3.75 × 10³ cells per well in 15 µl of culture medium in black-walled, clear-bottom 384-well plates (Greiner #781091) for forward pseudotyping. Cells were inoculated with 30 µl (96 well) or 5 ul (384 well) of tenfold-concentrated VSV pseudotyped virus and incubated for 16 h at 37 °C. VSV forward pseudotyping infection was assisted with spinfection at 1200 xg for 45 min a 4 °C. Cells were then lysed with 50 µl (96 well) or 15 µl (384 well) of 1× Renilla lysis buffer (Promega), prepared by diluting 5× Renilla lysis buffer in 1× PBS, and incubated for 20 min at room temperature with shaking.

Renilla luciferase substrate was prepared by adding 10 µl of substrate per milliliter of Renilla assay buffer (Promega), and 50 µl (96 well) or 15 µl (384 well) of the resulting reagent was added to each well. Luminescence was measured using a BioTek Synergy microplate reader and Gen5 software. Each condition was tested in at least technical triplicate within each independent experiment. Relative light units (RLU) were analyzed and plotted using GraphPad Prism v10.2.

### Lenti-GFP-EPCR entry into parahenipavirus G/F-expressing cells

Lentiviral GFP reporter particles bearing human EPCR (Lenti-GFP-EPCR) were generated to assess EPCR- dependent entry into cells expressing parahenipavirus G and F proteins. HEK-293T cells were co-transfected in 10-cm dishes with 6 µg of lentiviral Gag-Pol plasmid(*47*), 6 µg of human EPCR expression plasmid or empty-vector control, and 6 µg of pWPI-GFP (Addgene plasmid #12254) using PEI, as previously described(*34*).

At 48 h post-transfection, supernatants containing Lenti-GFP-EPCR particles were collected, passed through a 0.45 µm syringe filter, and concentrated tenfold using Amicon Ultra centrifugal filters with a 100 kDa MWCO.

For entry assays, 6 × 10⁵ HEK-293T cells were transfected in six-well plates with 1 µg each of the indicated parahenipavirus G and F expression plasmids using PEI. At 24 h post-transfection, 6 × 10⁴ G/F-expressing cells were seeded per well in 96-well plates and inoculated with 30 µl of concentrated Lenti-GFP-EPCR particles. Cells were incubated for 48 h at 37 °C, after which GFP-positive cells were quantified by flow cytometry using a Beckman Coulter CytoFLEX S. Flow cytometry data were analyzed using FlowJo v10, and entry was quantified as the percentage of GFP-positive cells.

### Protein C and anti-EPCR antibody inhibition assays

To assess inhibition of EPCR-dependent pseudotyped virus entry, 6 × 10⁵ HEK-293T cells were transfected with plasmids encoding the indicated parahenipavirus G and F proteins using PEI as described above. At 24 h post-transfection, 6 × 10⁴ cells were seeded per well in 96-well plates in 100 µl of DMEM.

Concentrated VSVΔG-luc-EPCR pseudotyped virus (30 µl) was incubated with 10 µg/ml anti-EPCR monoclonal antibody RCR-401 (BioLegend, #351902), 20 µg/ml His-tagged human protein C (Sino Biological, #10196-H08H), or PBS (Gibco, #14190144) as a negative control for 40 min at 4 °C. Virus–inhibitor mixtures were subsequently added to the cells and incubated for 16 h at 37 °C. Cells were lysed and Renilla luciferase activity was measured as described above.

### Meso Scale Discovery Binding Assay

Langya tetramer, Wenzhou tetramer, Mojiang tetramer, or Nipah trimer G proteins were coated on Meso Scale Discovery (MSD) 384-well bare plates (#L21XA) at 5 µg/mL overnight at 4°C. Plates were then washed and blocked with MSD Blocker A (#R93BA) for 1 hour at room temperature. After washing, serial dilutions of Sulfo-Tag labeled human EPCR Fc (Acro # PRR-H5254) from 0 to 10 µg/mL, in duplicate, were added to the plate for 1 hour at room temperature. Following incubation, plates were washed and read on an MSD Sector S600. Binding is reported as a function of electrochemiluminescence (ECL) units.

### Surface Plasmon Resonance

The binding affinity of human EPCR to Langya and Wenzhou tetramer G proteins was determined using an Alto high-throughput digital SPR system (Nicoya Lifesciences). Recombinant G proteins were immobilized on a Nicoya KC-CBX-CMD-16 sensor cartridge (3000-5000 response units) by amine coupling at 100 nM. Binding interactions to recombinant human EPCR Fc (ACRO # PRR-H5254) were evaluated by single-cycle kinetics in a running buffer of 10mM HEPES Buffer (10 mM HEPES, 150 mM NaCl, 3 mM EDTA, 0.1% Tween 20). Association was performed using 5 by 3-fold serial dilutions. Association occurred over 180 seconds, and dissociation occurred over 600 seconds. Specific SPR responses were calculated by subtracting reference sensor values from active sensor signals. Binding curves were fitted to a 1:1 Langmuir model. Calculated apparent constants are the average values of two independent experiments with an χ2/Rmax < 10%.

### Vaccination of nonhuman primates

All experiments were performed in accordance with ethical regulations at the respective institutions (Animal Care and Use Committees of the Vaccine Research Center, NIAID, NIH and of Bioqual, Inc). Macaques were housed and cared for in accordance with local, state, federal, and institute policies in facilities accredited by the American Association for Accreditation of Laboratory Animal Care (AAALAC), under standards established in the Animal Welfare Act and the Guide for the Care and Use of Laboratory Animals. Three rhesus macaques were immunized intramuscularly (IM) with a co-formulation of 100 mg of Langya G monomer (Genbank UUV47213.1, Asp 165 -Ser 624) and 100 mg of Nipah G monomer (Genbank AAK29088.1, Arg172-Thr 602) in 0.8 mL PBS. Just prior to IM administration, 20 % v/v Adjuplex (Cat # A0362, Millipore Sigma) was added for a final volume of 1 mL. The dose was split and delivered to the right and left quadriceps. Animals were boosted at weeks 4, 16 and 24 with the same formulation and procedure as the prime. Two weeks after each immunization, blood was collected and used to isolate plasma for neutralization assays. At the end of the protocol, the animals were retained for future studies.

### Nonhuman primate (NHP) plasma dilution assays

NHP plasma was aliquoted such that samples were subject to no more than 3 freeze-thaws. Plasma isolated from NHPs at 18 weeks was serially diluted by half log_10_ factors in DMEM supplemented with 2% FBS in 96-well plates. 11 µl of tenfold-concentrated VSV pseudotyped virus was added to 16 µl of diluted plasma. BHK cells stably expressing human EPCR cells were seeded at 3.75 × 10³ cells per well in 15 µl of culture medium in black-walled, clear bottom 384-well plates (Greiner #781091). Cells were inoculated with 10 µl of diluted plasma-virus mix and the remainder of the protocol is as written above for VSV pseudotyping entry assay.

### EPCR receptor orthologs

EPCR receptor ortholog sequences were manually extracted from NCBI with accession numbers indicated in Data S4. *Crocidura gueldenstaedtii* and *Blarina brevicauda* did not have an annotated EPCR sequence. We applied the miniprot algorithm(*48*) to the *Crocidura gueldenstaedtii* (GCA_051295465.2) and *Blarina brevicauda* (GCA_965212715.2) genomes using the *Sorex araneus* annotated EPCR (XP_004612598.1) as the query and using the splice aware option. The extracted sequences are included in Data S4. The *Rattus tanezumi*, formerly *Rattus flavipectus*, genome assembly and EPCR sequence was also not available.

Publicly available *Rattus tanezumi* raw sequences (SRR3948161 and SRR2917427) were parsed against the EPCR sequence of Rattus rattus +/- a 25 kbp region and the matching reads were then mapped with minimap2(*49*) and consensus constructed with samtools(*50*) based on the *Rattus rattus* reference genome (GCF_011064425.1). After reference-based genome construction, the *Rattus tanezumi* protein sequence was derived using the miniprot algorithm with the *Rattus norvegicus* EPCR sequence (NP_001020904.1) as query.

### Cryo-EM grid preparation, data collection and structure determination

The complex of Langya G and human endothelial protein C receptor (EPCR) was formed by mixing Langya G and EPCR to a final concentration of 0.19 mg/mL for each component. The complex was incubated for 10 min at room temperature. Before vitrification, n-Dodecyl-β-D-maltoside (DDM) was added to the sample to a final concentration of 0.085 mM. A 2.7-µL drop of the complex was applied to glow-discharged Quantifoil R 1.2/1.3 gold grids and plunge-frozen using a Vitrobot Mark IV with blot times ranging from 1.5 to 2 s (95% humidity, 4 °C, blot force of -5).

Cryo-EM datasets were collected on a Titan Krios G1 microscope equipped with a Gatan K3 detector and a Gatan BioContinuum Imaging Filter using SerialEM(*51*). In total, 5,174 movies at a 35° of stage tilt and 6,464 movies at a 45° tilt were recorded at a 105,000x nominal magnification in the super-resolution mode, corresponding to a pixel size of 0.4183 Å. Motion correction and CTF parameter estimation were performed in cryoSPARC v5.0.6(*52*) using Patch Motion Correction and Patch CTF estimation. After curation to remove micrographs with low CTF-estimated resolution or thick ice, 10,580 micrographs were retained for downstream data processing.

Particles were initially picked automatically with crYOLO from a subset of 500 micrographs(*53*). After several rounds of 2D classification, high-resolution classes representing different particle orientations were selected as templates for template-based particle picking and Topaz training(*54*). Particles from template-based picking and Topaz picking were merged and duplicates were removed. Ab initio model generation was followed by several rounds of heterogeneous refinement in cryoSPARC, yielding 182,884 high-quality particles that were used for the final non-uniform refinement with C4 symmetry imposed. The final reconstruction of the Langya G-EPCR complex reached a resolution of 3.1 Å (**Table S1, Figure S2**) as determined using the gold-standard FSC 0.143 criterion^35^.

Initial atomic models of Langya G (PDB 8JZB) and EPCR (PDB 1L8J(*28*)) were docked into the cryo-EM map using UCSF ChimeraX(*55*) and refined in Coot(*56*) and ISOLDE(*57*) using the sharpened map generated by cryoSPARC and a post-processed map generated with DeepEMhancer(*58*). The final model was refined in Phenix(*59*) using real-space refinement against the original cryoSPARC map. Model quality was validated using Molprobity(*60*) (**Table S1**). Protein-protein interfaces were analyzed using PISA(*61*), and figures were prepared using ChimeraX.

The interface between Langya G glycoprotein and hEPCR was calculated using QtPISA (CCP4)(*61*). Hydrogen bonds and salt bridges were assigned using the criteria implemented in PISA. (**Table S2**)

### Statistical analysis

Pseudovirus experiments were performed two to four independent times with three technical replicates per experiment. IC_50_ values were determined by non-linear regression. Data were analyzed in GraphPad Prism (v11.1.0) and are presented as mean ± standard deviation. Statistical tests are stated in the figure legends. *P*<0.05 was considered significant; \**P*<0.05, \*\**P*<0.01, \*\*\**P*<0.001, \*\*\*\**P*<0.0001.

### Data, code, and materials availability

Cryo-EM structural data of the Langya G in complex with EPCR generated here has been deposited to the Protein Data Bank (PDB) under accession number 39FI and cryo-EM maps have been deposited to the Electron Microscopy Data Bank with accession code EMDB-79384. All data are available in the main text or supplementary materials. Code used to generate figure 1A, analyze CRISPRa data, and extract EPCR sequences are available at github repository: https://github.com/pmtran5884/Langya_paper/. All materials are available to qualified investigators with a materials transfer agreement (MTA) and appropriate biosafety approval.

## Author contributions

Conceptualization: LS, AS, CBW, TZ, DCD, ZZ, PT

Methodology: NL, LO, JS, CBW, ZZ, PT, TZ, LS

Investigation: ZZ, PT, NL, LO, MS, MC, AC, JAB, JMW, TTD, BZ, JZ, ACP, CM, AS, DRM, WM, RP, AS, SL, YT, LS, TZ

Project management: AS

Visualization: ZZ, PT, CBW, TZ, YT, NL, LI, LS

Funding acquisition: CBW, TZ, WM, RP, SL, AS, DCD, LS;

Writing—original draft: CBW, TZ, LS, ZZ, PT

Reviewing and editing: all authors.

## Funding

This research was supported in part by the Intramural Research Program of the National Institutes of Health (NIH). The contributions of the NIH author(s) are considered Works of the United States Government. The findings and conclusions presented in this paper are those of the author(s) and do not necessarily reflect the views of the NIH or the U.S. Department of Health and Human Services. This work was funded in part by the Frederick National Laboratory for Cancer Research, NIH, under contract 75N91019D00024 and National Institutes of Health, Intramural Research Program of the Vaccine Research Center and the Division of Intramural Research. Sequencing was performed at YCGA which was supported by the National Institute of General Medical Sciences of the National Institutes of Health under Award Number 1S10OD030363-01A1. CBW was supported by Burroughs Wellcome Fund and DARPA YFA.

## Competing interests

Authors declare no competing interests.

## Acknowledgements

We thank Robert Orchard, Arthur Kim, Arya Ökten, and Charlotte Shore for helpful discussions. We thank Christopher Broder for HeLa-USU cells. We also thank Forest Hoyt and Vinod Nair of the Rocky Mountain Laboratories, NIAID, NIH, for cryo-EM data collection. This research was supported in part by the Vaccine Research Center of the National Institute of Allergy and Infectious Diseases, National Institutes of Health (NIH). The contributions of the NIH authors are considered Works of the United States Government. The findings and conclusions presented in this paper are those of the authors and do not necessarily reflect the views of the NIH or the U.S. Department of Health and Human Services.

## Notes

### Competing Interest Statement

The authors have declared no competing interest.

## References

1. K. B. Chua, W. J. Bellini, P. A. Rota, B. H. Harcourt, A. Tamin, S. K. Lam, T. G. Ksiazek, P. E. Rollin, S. R. Zaki, W. Shieh, C. S. Goldsmith, D. J. Gubler, J. T. Roehrig, B. Eaton, A. R. Gould, J. Olson, H. Field, P. Daniels, A. E. Ling, C. J. Peters, L. J. Anderson, B. W. Mahy, Nipah virus: a recently emergent deadly paramyxovirus. Science 288, 1432–1435 (2000).

2. K. Murray, P. Selleck, P. Hooper, A. Hyatt, A. Gould, L. Gleeson, H. Westbury, L. Hiley, L. Selvey, B. Rodwell, A morbillivirus that caused fatal disease in horses and humans. Science 268, 94–97 (1995).

3. X.-A. Zhang, H. Li, F.-C. Jiang, F. Zhu, Y.-F. Zhang, J.-J. Chen, C.-W. Tan, D. E. Anderson, H. Fan, L.-Y. Dong, C. Li, P.-H. Zhang, Y. Li, H. Ding, L.-Q. Fang, L.-F. Wang, W. Liu, A Zoonotic Henipavirus in Febrile Patients in China. N. Engl. J. Med. 387, 470–472 (2022).

4. Y.-M. Chen, S.-J. Hu, X.-D. Lin, J.-H. Tian, J.-X. Lv, M.-R. Wang, X.-Q. Luo, Y.-Y. Pei, R.-X. Hu, Z.-G. Song, E. C. Holmes, Y.-Z. Zhang, Host traits shape virome composition and virus transmission in wild small mammals. Cell 186, 4662–4675.e12 (2023).

5. Z. Wu, L. Yang, F. Yang, X. Ren, J. Jiang, J. Dong, L. Sun, Y. Zhu, H. Zhou, Q. Jin, Novel Henipa-like virus, Mojiang Paramyxovirus, in rats, China, 2012. Emerg. Infect. Dis. 20, 1064–1066 (2014).

6. L. Shen, Y. Tong, W. Chai, J. Sun, J. Zhao, L. Yin, Y. Yang, C. Wang, W. Tan, J. Zhang, Parahenipavirus langyaense (Langya virus): Spillover ecology, noncanonical receptor-mediated entry, and future control strategies. One Health 23, 101521 (2026).

7. R. H. Parry, K. Y. H. Yamada, W. R. Hood, Y. Zhao, J. Y. Lu, A. Seluanov, V. Gorbunova, N. Modhiran, D. Watterson, A. Isaacs, Henipavirus in Northern Short-Tailed Shrew, Alabama, USA. Emerg. Infect. Dis. 31, 392–394 (2025).

8. S. Madera, A. Kistler, H. C. Ranaivoson, V. Ahyong, A. Andrianiaina, S. Andry, V. Raharinosy, T. H. Randriambolamanantsoa, N. A. F. Ravelomanantsoa, C. M. Tato, J. L. DeRisi, H. C. Aguilar, V. Lacoste, P. Dussart, J.-M. Heraud, C. E. Brook, Discovery and Genomic Characterization of a Novel Henipavirus, Angavokely Virus, from Fruit Bats in Madagascar. J. Virol. 96, e0092122 (2022).

9. K. Xu, K. R. Rajashankar, Y.-P. Chan, J. P. Himanen, C. C. Broder, D. B. Nikolov, Host cell recognition by the henipaviruses: crystal structures of the Nipah G attachment glycoprotein and its complex with ephrin-B3. Proc. Natl. Acad. Sci. U. S. A. 105, 9953–9958 (2008).

10. Z. Wang, M. Amaya, A. Addetia, H. V. Dang, G. Reggiano, L. Yan, A. C. Hickey, F. DiMaio, C. C. Broder, D. Veesler, Architecture and antigenicity of the Nipah virus attachment glycoprotein. Science 375, 1373–1378 (2022).

11. C. T. Pager, R. E. Dutch, Cathepsin L is involved in proteolytic processing of the Hendra virus fusion protein. J. Virol. 79, 12714–12720 (2005).

12. S. Diederich, L. Sauerhering, M. Weis, H. Altmeppen, N. Schaschke, T. Reinheckel, S. Erbar, A. Maisner, Activation of the Nipah virus fusion protein in MDCK cells is mediated by cathepsin B within the endosome-recycling compartment. J. Virol. 86, 3736–3745 (2012).

13. O. A. Negrete, E. L. Levroney, H. C. Aguilar, A. Bertolotti-Ciarlet, R. Nazarian, S. Tajyar, B. Lee, EphrinB2 is the entry receptor for Nipah virus, an emergent deadly paramyxovirus. Nature 436, 401–405 (2005).

14. M. I. Bonaparte, A. S. Dimitrov, K. N. Bossart, G. Crameri, B. A. Mungall, K. A. Bishop, V. Choudhry, D. S. Dimitrov, L.-F. Wang, B. T. Eaton, C. C. Broder, Ephrin-B2 ligand is a functional receptor for Hendra virus and Nipah virus. Proc. Natl. Acad. Sci. U. S. A. 102, 10652–10657 (2005).

15. S. Cheliout Da Silva, L. Yan, H. V. Dang, K. Xu, J. H. Epstein, D. Veesler, C. C. Broder, Functional Analysis of the Fusion and Attachment Glycoproteins of Mojiang Henipavirus. Viruses 13, 517 (2021).

16. O. A. Negrete, M. C. Wolf, H. C. Aguilar, S. Enterlein, W. Wang, E. Mühlberger, S. V. Su, A. Bertolotti-Ciarlet, R. Flick, B. Lee, Two key residues in ephrinB3 are critical for its use as an alternative receptor for Nipah virus. PLoS Pathog. 2, e7 (2006).

17. A. Natasha, S. E. Pye, K. Park, S. Rajoriya, I. Yang, J. Park, H. S. Pangestu, J. Kim, Y. Oh, C. B. López, J.-W. Song, W.-K. Kim, Detection and characterization of Langya virus in Crocidura lasiura (the Ussuri white-toothed shrew), Republic of Korea. One Health Amst. Neth. 20, 101017 (2025).

18. M. I. Bonaparte, A. S. Dimitrov, K. N. Bossart, G. Crameri, B. A. Mungall, K. A. Bishop, V. Choudhry, D. S. Dimitrov, L.-F. Wang, B. T. Eaton, C. C. Broder, Ephrin-B2 ligand is a functional receptor for Hendra virus and Nipah virus. Proc. Natl. Acad. Sci. U. S. A. 102, 10652–10657 (2005).

19. S. Madera, A. Kistler, H. C. Ranaivoson, V. Ahyong, A. Andrianiaina, S. Andry, V. Raharinosy, T. H. Randriambolamanantsoa, N. A. F. Ravelomanantsoa, C. M. Tato, J. L. DeRisi, H. C. Aguilar, V. Lacoste, P. Dussart, J.-M. Heraud, C. E. Brook, Discovery and Genomic Characterization of a Novel Henipavirus, Angavokely Virus, from Fruit Bats in Madagascar. J. Virol. 96, e0092122 (2022).

20. S.-H. Lee, K. Kim, J. Kim, J. S. No, K. Park, S. Budhathoki, S. H. Lee, J. Lee, S. H. Cho, S. Cho, G.-Y. Lee, J. Hwang, H.-C. Kim, T. A. Klein, C.-S. Uhm, W.-K. Kim, J.-W. Song, Discovery and Genetic Characterization of Novel Paramyxoviruses Related to the Genus Henipavirus in Crocidura Species in the Republic of Korea. Viruses 13, 2020 (2021).

21. B. Vanmechelen, S. Meurs, M. Horemans, A. Loosen, T. Joly Maes, L. Laenen, V. Vergote, F. R. Koundouno, N. Magassouba, M. K. Konde, I. S. Condé, M. W. Carroll, P. Maes, The characterization of multiple novel paramyxoviruses highlights the diverse nature of the subfamily Orthoparamyxovirinae. Virus Evol. 8, veac061 (2022).

22. M. Horemans, J. Van Bets, T. Joly Maes, P. Maes, B. Vanmechelen, Discovery and genome characterization of six new orthoparamyxoviruses in small Belgian mammals. Virus Evol. 9, vead065 (2023).

23. J. Barr, S. Caruso, S. J. Edwards, S. Todd, I. Smith, M. Tachedjian, G. Crameri, L.-F. Wang, G. A. Marsh, Novel Henipavirus, Salt Gully Virus, Isolated from Pteropid Bats, Australia. Emerg. Infect. Dis. 31, 1824–1828 (2025).

24. J. Song, R. D. Chow, M. A. Peña-Hernández, L. Zhang, S. A. Loeb, E.-Y. So, O. D. Liang, P. Ren, S. Chen, C. B. Wilen, S. Lee, LRRC15 inhibits SARS-CoV-2 cellular entry in trans. PLoS Biol. 20, e3001805 (2022).

25. L. V. Mohan Rao, C. T. Esmon, U. R. Pendurthi, Endothelial cell protein C receptor: a multiliganded and multifunctional receptor. Blood 124, 1553–1562 (2014).

26. K. Fukudome, C. T. Esmon, Identification, cloning, and regulation of a novel endothelial cell protein C/activated protein C receptor. J. Biol. Chem. 269, 26486–26491 (1994).

27. E. Erausquin, M. Morán-Garrido, J. Sáiz, C. Barbas, G. Dichiara-Rodríguez, A. Urdiciain, J. López-Sagaseta, Identification of a broad lipid repertoire associated to the endothelial cell protein C receptor (EPCR). Sci. Rep. 12, 15127 (2022).

28. V. Oganesyan, N. Oganesyan, S. Terzyan, D. Qu, Z. Dauter, N. L. Esmon, C. T. Esmon, The crystal structure of the endothelial protein C receptor and a bound phospholipid. J. Biol. Chem. 277, 24851–24854 (2002).

29. M. Riewald, R. J. Petrovan, A. Donner, B. M. Mueller, W. Ruf, Activation of endothelial cell protease activated receptor 1 by the protein C pathway. Science 296, 1880–1882 (2002).

30. L. Turner, T. Lavstsen, S. S. Berger, C. W. Wang, J. E. V. Petersen, M. Avril, A. J. Brazier, J. Freeth, J. S. Jespersen, M. A. Nielsen, P. Magistrado, J. Lusingu, J. D. Smith, M. K. Higgins, T. G. Theander, Severe malaria is associated with parasite binding to endothelial protein C receptor. Nature 498, 502–505 (2013).

31. C. K. Y. Lau, L. Turner, J. S. Jespersen, E. D. Lowe, B. Petersen, C. W. Wang, J. E. V. Petersen, J. Lusingu, T. G. Theander, T. Lavstsen, M. K. Higgins, Structural conservation despite huge sequence diversity allows EPCR binding by the PfEMP1 family implicated in severe childhood malaria. Cell Host Microbe 17, 118–129 (2015).

32. G. D. Haas, O. Escaffre, R. A. Reis, T. L. Juelich, J. K. Smith, L. Zhang, B. K. Kalveram, A. A. Guzmán-Solís, D. Vyshenska, W. Klain, A. L. Greninger, A. N. Freiberg, B. Lee, De novo recovery of Ghana virus, an African bat Henipavirus, reveals differential tropism and attenuated pathogenicity compared to Nipah virus. Cell Rep. 45, 117074 (2026).

33. B. Lee, O. Pernet, A. A. Ahmed, A. Zeltina, S. M. Beaty, T. A. Bowden, Molecular recognition of human ephrinB2 cell surface receptor by an emergent African henipavirus. Proc. Natl. Acad. Sci. U. S. A. 112, E2156–2165 (2015).

34. M. Madel Alfajaro, E. L. Keeler, N. Li, N. J. Catanzaro, I.-T. Teng, Z. Zhao, M. W. Grunst, B. Yount, A. Schäfer, D. Wang, A. S. Kim, A. Synowiec, M. A. Peña-Hernández, S. Zepeda, R. Arinola, R. Kaur, B. L. Menasche, J. Wei, G. A. Russell, J. Huck, J. Song, A. Ring, A. Iwasaki, R. K. Jangra, S. Lee, D. R. Martinez, W. Mothes, P. D. Uchil, J. G. Doench, A. B. Spaulding, R. S. Baric, L. Serebryannyy, Y. Tsybovsky, T. Zhou, D. C. Douek, C. B. Wilen, HKU5 bat merbecoviruses engage bat and mink ACE2 as entry receptors. Nat. Commun. 16, 6822 (2025).

35. W. Li, S. Wu, Q. Gui, C. Liu, B. Zhou, H. Yan, Y. Sun, Q. Fan, Y. Zhou, H. Guo, S. Tang, X. Ge, B. Ju, R. Yan, Z. Zhang, Structure and function of a pair of non-competing monoclonal antibodies against Langya henipavirus attachment glycoprotein. Cell Rep. 44, 116407 (2025).

36. T. A. Bowden, A. R. Aricescu, R. J. C. Gilbert, J. M. Grimes, E. Y. Jones, D. I. Stuart, Structural basis of Nipah and Hendra virus attachment to their cell-surface receptor ephrin-B2. Nat. Struct. Mol. Biol. 15, 567–572 (2008).

37. E. D. Laing, C. K. Navaratnarajah, S. Cheliout Da Silva, S. R. Petzing, Y. Xu, S. L. Sterling, G. A. Marsh, L.-F. Wang, M. Amaya, D. B. Nikolov, R. Cattaneo, C. C. Broder, K. Xu, Structural and functional analyses reveal promiscuous and species specific use of ephrin receptors by Cedar virus. Proc. Natl. Acad. Sci. U. S. A. 116, 20707–20715 (2019).

38. E. D. Laing, M. Amaya, C. K. Navaratnarajah, Y.-R. Feng, R. Cattaneo, L.-F. Wang, C. C. Broder, Rescue and characterization of recombinant cedar virus, a non-pathogenic Henipavirus species. Virol. J. 15, 56 (2018).

39. M. Pagès, Y. Chaval, V. Herbreteau, S. Waengsothorn, J.-F. Cosson, J.-P. Hugot, S. Morand, J. Michaux, Revisiting the taxonomy of the Rattini tribe: a phylogeny-based delimitation of species boundaries. BMC Evol. Biol. 10, 184 (2010).

40. J.-M. Gu, J. T. B. Crawley, G. Ferrell, F. Zhang, W. Li, N. L. Esmon, C. T. Esmon, Disruption of the endothelial cell protein C receptor gene in mice causes placental thrombosis and early embryonic lethality. J. Biol. Chem. 277, 43335–43343 (2002).

41. A. Kucukelbir, F. J. Sigworth, H. D. Tagare, Quantifying the local resolution of cryo-EM density maps. Nat. Methods 11, 63–65 (2014).

42. K. Katoh, K. Misawa, K. Kuma, T. Miyata, MAFFT: a novel method for rapid multiple sequence alignment based on fast Fourier transform. Nucleic Acids Res. 30, 3059–3066 (2002).

43. S. Kumar, G. Stecher, M. Suleski, M. Sanderford, S. Sharma, K. Tamura, MEGA12: Molecular Evolutionary Genetic Analysis Version 12 for Adaptive and Green Computing. Mol. Biol. Evol. 41, msae263 (2024).

44. S. Konermann, M. D. Brigham, A. E. Trevino, J. Joung, O. O. Abudayyeh, C. Barcena, P. D. Hsu, N. Habib, J. S. Gootenberg, H. Nishimasu, O. Nureki, F. Zhang, Genome-scale transcriptional activation by an engineered CRISPR-Cas9 complex. Nature 517, 583–588 (2015).

45. K. R. Sanson, R. E. Hanna, M. Hegde, K. F. Donovan, C. Strand, M. E. Sullender, E. W. Vaimberg, A. Goodale, D. E. Root, F. Piccioni, J. G. Doench, Optimized libraries for CRISPR-Cas9 genetic screens with multiple modalities. Nat. Commun. 9, 5416 (2018).

46. M. S. Strine, W. L. Cai, J. Wei, M. M. Alfajaro, R. B. Filler, S. B. Biering, S. Sarnik, R. D. Chow, A. Patil, K. S. Cervantes, C. K. Collings, P. C. DeWeirdt, R. E. Hanna, K. Schofield, C. Hulme, S. Konermann, J. G. Doench, P. D. Hsu, C. Kadoch, Q. Yan, C. B. Wilen, DYRK1A promotes viral entry of highly pathogenic human coronaviruses in a kinase-independent manner. PLoS Biol. 21, e3002097 (2023).

47. Z. Zhao, R. Fagerlund, H. Tossavainen, K. Hopfensperger, R. Lotke, S. Srinivasachar Badarinarayan, F. Kirchhoff, P. Permi, K. Sato, D. Sauter, K. Saksela, Evolutionary plasticity of SH3 domain binding by Nef proteins of the HIV-1/SIVcpz lentiviral lineage. PLoS Pathog. 17, e1009728 (2021).

48. H. Li, Protein-to-genome alignment with miniprot. Bioinformatics 39, btad014 (2023).

49. H. Li, Minimap2: pairwise alignment for nucleotide sequences. Bioinformatics 34, 3094– 3100 (2018).

50. H. Li, B. Handsaker, A. Wysoker, T. Fennell, J. Ruan, N. Homer, G. Marth, G. Abecasis, R. Durbin, 1000 Genome Project Data Processing Subgroup, The Sequence Alignment/Map format and SAMtools. Bioinformatics 25, 2078–2079 (2009).

51. D. N. Mastronarde, Automated electron microscope tomography using robust prediction of specimen movements. J. Struct. Biol. 152, 36–51 (2005).

52. A. Punjani, J. L. Rubinstein, D. J. Fleet, M. A. Brubaker, cryoSPARC: algorithms for rapid unsupervised cryo-EM structure determination. Nat. Methods 14, 290–296 (2017).

53. T. Wagner, F. Merino, M. Stabrin, T. Moriya, C. Antoni, A. Apelbaum, P. Hagel, O. Sitsel, T. Raisch, D. Prumbaum, D. Quentin, D. Roderer, S. Tacke, B. Siebolds, E. Schubert, T. R. Shaikh, P. Lill, C. Gatsogiannis, S. Raunser, SPHIRE-crYOLO is a fast and accurate fully automated particle picker for cryo-EM. *Commun*. Biol. 2, 218 (2019).

54. T. Bepler, A. Morin, M. Rapp, J. Brasch, L. Shapiro, A. J. Noble, B. Berger, Positive-unlabeled convolutional neural networks for particle picking in cryo-electron micrographs. Nat. Methods 16, 1153–1160 (2019).

55. E. C. Meng, T. D. Goddard, E. F. Pettersen, G. S. Couch, Z. J. Pearson, J. H. Morris, T. E. Ferrin, UCSF ChimeraX: Tools for structure building and analysis. Protein Sci. Publ. Protein Soc. 32, e4792 (2023).

56. P. Emsley, K. Cowtan, Coot: model-building tools for molecular graphics. Acta Crystallogr. D Biol. Crystallogr. 60, 2126–2132 (2004).

57. T. I. Croll, ISOLDE: a physically realistic environment for model building into low-resolution electron-density maps. Acta Crystallogr. Sect. Struct. Biol. 74, 519–530 (2018).

58. R. Sanchez-Garcia, J. Gomez-Blanco, A. Cuervo, J. M. Carazo, C. O. S. Sorzano, J. Vargas, DeepEMhancer: a deep learning solution for cryo-EM volume post-processing. *Commun*. Biol. 4, 874 (2021).

59. D. Liebschner, P. V. Afonine, M. L. Baker, G. Bunkóczi, V. B. Chen, T. I. Croll, B. Hintze, L. W. Hung, S. Jain, A. J. McCoy, N. W. Moriarty, R. D. Oeffner, B. K. Poon, M. G. Prisant, R. J. Read, J. S. Richardson, D. C. Richardson, M. D. Sammito, O. V. Sobolev, D. H. Stockwell, T. C. Terwilliger, A. G. Urzhumtsev, L. L. Videau, C. J. Williams, P. D. Adams, Macromolecular structure determination using X-rays, neutrons and electrons: recent developments in Phenix. Acta Crystallogr. Sect. Struct. Biol. 75, 861–877 (2019).

60. I. W. Davis, L. W. Murray, J. S. Richardson, D. C. Richardson, MOLPROBITY: structure validation and all-atom contact analysis for nucleic acids and their complexes. Nucleic Acids Res. 32, W615–619 (2004).

61. E. Krissinel, K. Henrick, Inference of macromolecular assemblies from crystalline state. J. Mol. Biol. 372, 774–797 (2007).

